# Tracking bacteria at high density with FAST, the Feature-Assisted Segmenter/Tracker

**DOI:** 10.1101/2021.11.26.470050

**Authors:** Oliver J. Meacock, William M. Durham

## Abstract

Most bacteria live attached to surfaces in densely-packed communities^1,2^. While new experimental and imaging techniques are beginning to provide a window on the complex processes that play out in these communities, resolving the behaviour of individual cells through time and space remains a major challenge. Although a number of different software solutions have been developed to track microorganisms^3–8^, these approaches typically rely on a large number of user-defined parameters that must be carefully tuned to effectively track cells. Testing a given parameter combination can take hours to days depending on the size of the dataset, making iterative optimisation impractical. To overcome these limitations, we have developed FAST, the Feature-Assisted Segmenter/Tracker, which uses unsupervised machine learning to optimise tracking while maintaining ease of use. Our approach, rooted in information theory, largely eliminates the need for users to iteratively adjust parameters manually and make qualitative assessments of the resulting cell trajectories. Instead, FAST measures multiple distinguishing “features” for each cell and then autonomously quantifies the amount of unique information each feature provides. We then use these measurements to determine how data from different features should be combined to minimize tracking errors. Comparing our algorithm with a naïve approach that uses cell position alone revealed that FAST produced 4 to 10 times fewer tracking errors. The modular design of FAST combines our novel tracking method with tools for segmentation, extensive data visualisation, lineage assignment, and manual track correction. It is also highly extensible, allowing users to extract custom information from images and seamlessly integrate it into downstream analyses. FAST therefore enables high-throughput, data-rich analyses with minimal user input. It has been released for use either in Matlab or as a compiled stand-alone application, and is available at https://bit.ly/3vovDHn, along with extensive tutorials and detailed documentation.

## Introduction

Time-lapse microscopic imaging and automated cell tracking has led to many fundamental advances in our understanding of how microorganisms sense and respond to their environment. While many studies have focused on the movement of planktonic bacteria at relatively low densities, many behaviours – including collective movement^9,10^, combat^11^, sharing of public goods^12^ and genetic exchange^13^ – typically only occur in the closely-packed assemblages in which most microbes live. These dense communities are often studied in the laboratory using confluent monolayers of cells, which are much easier to image than three-dimensional aggregations. One method to generate such monolayers is to confine cells with a slab of agarose or polyacrylamide to form an interstitial colony^9,14–16^, while more advanced microfluidic techniques^17^ can also be used to confine cells to a single plane while allowing for more precise control over their chemical environment (Fig. 1a). Monolayers can also form in thin films of fluid, including those arising naturally during bacterial swarming motility^10^ and in assays used to study mixing induced by flagellar motility^18^. Regardless of the origin of the monolayer however, investigators face the same technical challenges when tracking densely packed cells using phase-contrast, brightfield and/or epi-fluorescence microscopy (Fig. 1b).

**Fig. 1.**
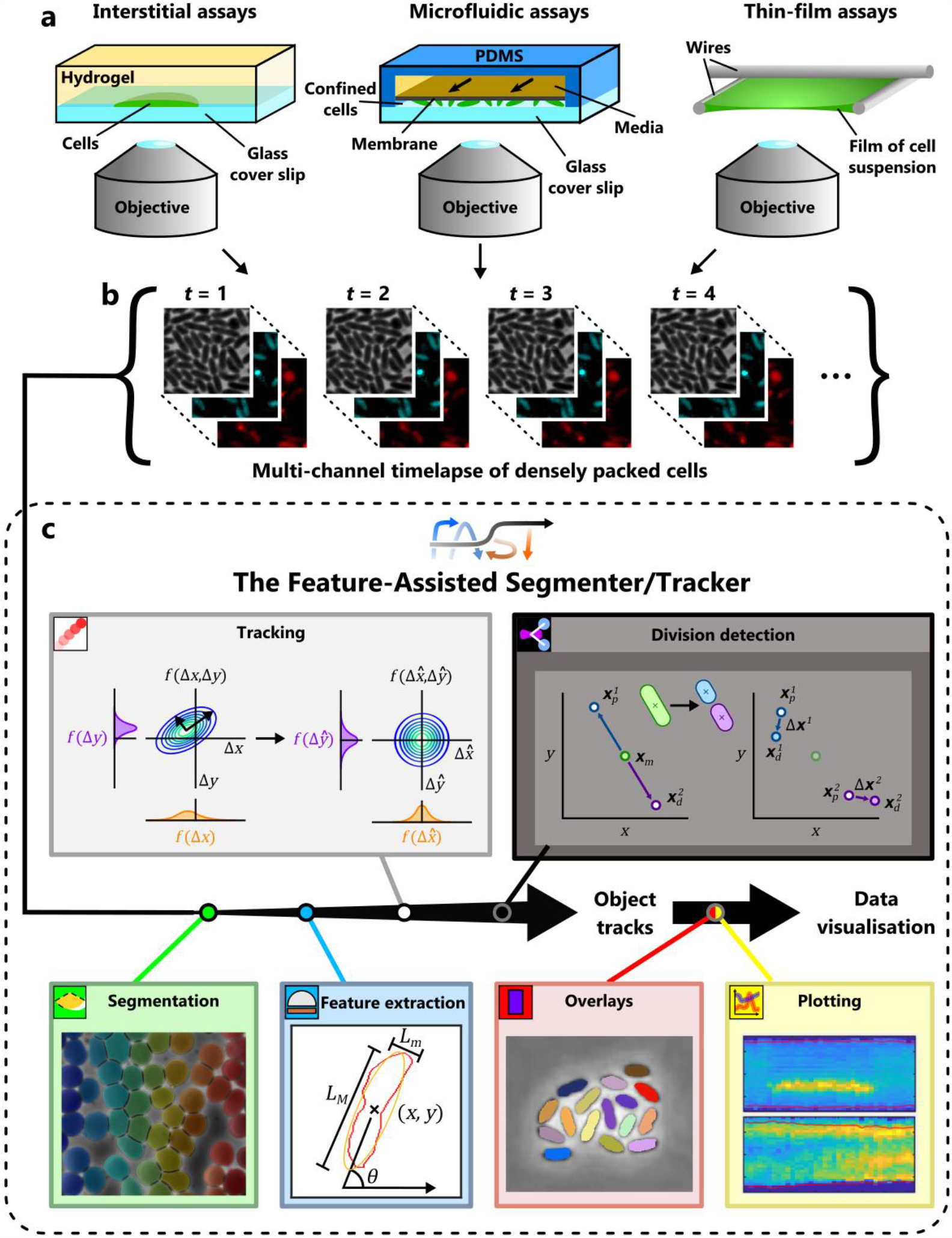
Analysis of high-density datasets using FAST’s modular framework. (a) While bacteria naturally form monolayers in some environments, a number of different assays are used to physically confine cells to a plane in laboratory experiments. Among these are interstitial colonies, microfluidic devices that trap cells between two solid boundaries, and experimental assays that confine cells to a thin film of liquid. (b) These experiments are typically imaged using automated microscopy systems capable of collecting images in both transmitted light and fluorescent channels at specified time points. (c) FAST analyses these imaging datasets in six separate modules, each of which is used in sequence (see text). A complete specification of each module is provided in the Supplementary Information.

Tracking solitary cells at low density is relatively straightforward, and basic tracking algorithms that use cell position alone often produce excellent results. However, tracking cells that are densely packed together is notoriously difficult because the spacing between neighbouring cells becomes similar to the distance cells move between frames. These problems are exacerbated by cell motility. While some species of bacteria are non-motile at high density and spread slowly only via cell division and the secretion of surfactants^15,17^, emergent patterns of collective motility driven by twitching^9^, swarming^10^ and gliding^19^ further complicate tracking by rapidly changing the positions of cells. Consequently, dense, motile communities must be imaged at high framerates for tracking to be feasible, such that a typical experiment requires a timeseries of hundreds or thousands of images. The large size of imaging datasets, combined with the large number of cells within each image, means that the computational time required for cell tracking is often one of the main bottlenecks in a researcher’s workflow. This can be compounded if tracking must be repeated multiple times to optimise tracking parameters.

A basic nearest-neighbour tracking algorithm compares the coordinates of cell centroids between subsequent frames and build trajectories by connecting those centroids that are closest together between subsequent time points. More advanced algorithms leverage additional cell characteristics or “features” to distinguish cells from one another, including metrics that measure cell shape, orientation, fluorescence levels and patterns of previous movement^3,20–22^. However, incorporating these additional feature measurements introduces a new problem, namely how to efficiently combine the information from different features to optimise tracking performance. In some software packages (e.g. MicrobeJ^3^), this requires the user to manually choose a large number of parameters and make qualitative judgements of the trajectories that result from each set of parameters. The combinatorial explosion in the number of possible parameters, the fact that a single parameter set can require many hours to test, and the lack of a rigorous way to compare the tracking results across different parameter sets, means that parameter optimisation in such a tracking algorithm is typically a highly iterative, time-consuming process.

A related problem is the fact that properties of bacteria within collectives are highly dynamic, changing at both the individual level and population level over time. For example, fluorophores can accumulate within cells or bleach, while cell movement can speed up or slow down due to the secretion of extracellular factors or changes in gene expression^23,24^. In addition, the overall density of cells often increases over time as a result of cell division. This variability means that an algorithm optimised to track cells at the beginning of an experiment might struggle at later timepoints. Furthermore, experimental issues - such as changes in illumination, focus, or shifts in the field of view caused by thermal drift - can cause an abrupt deterioration of tracking accuracy. Knowing when tracking accuracy has deteriorated to an unacceptable level often lacks a rigorous basis and requires the output of tracking software to be carefully validated by eye, typically infeasible for high-throughput datasets.

In this paper, we discuss a new approach that uses unsupervised machine learning to improve the fidelity of tracking. Our system automatically measures the statistical properties of each feature over time and then uses this data to dynamically change the relative weighting of each feature based on the information it can contribute to solving the tracking problem. It also provides users with a metric of the expected accuracy of the resulting cell trajectories, alerting them to sections of datasets that may need to be omitted from subsequent analyses. This tracking algorithm is combined with robust segmentation, feature extraction, lineage analysis and visualisation routines to make up FAST, the Feature-Assisted Segmenter/Tracker. FAST has been released as open-source software which can be run either directly within Matlab or as a stand-alone application, and has already been used in a number of publications to accurately analyse densely-packed bacterial monolayers^9,16,25,26^.

In the following sections, we discuss the design approach and general structure of FAST, and then illustrate the utility of our novel cell tracking approach using synthetic datasets. Next, we discuss three case studies that illustrate the versatility of FAST, including: 1) tracking of twitching *P. aeruginosa* cells in a 2D monolayer, 2) lineage analysis of an *E. coli* colony, and 3) automated analysis of the Type 6 Secretion System (T6SS) in a co-culture of *P. aeruginosa* and *V. cholerae*. While these case studies focus on densely packed bacteria, FAST can also be used to analyse other types of biological samples (*e*.*g*. Fig. S1g-i).

## Results

### Software overview

Initially, we conducted a review of existing cell tracking software packages^3–8^ to establish four key design objectives for our software: modularity, rapid user feedbacks, minimisation of user-defined parameters and extensibility (see section 2 of the Supplementary Information for further details). We built the FAST pipeline following these design principles, resulting in a set of six modules that are used in sequence (Fig. 1c). First, the Segmentation module is used to identify the boundaries of individual cells, typically using brightfield or phase-contrast images. Next, the Feature Extraction module measures a range of different cell properties such as position, size and fluorescence intensity (potentially in multiple channels) using the previously extracted segmentation as a basis. The Tracking module employs our machine learning process to quantify the information associated with each feature and then calculates the relative weighting of each feature to maximise tracking fidelity. If required, a manual validation and correction sub-module also allows the user to correct any mistakes made by the tracking algorithm. The optional Division Detection module uses a closely related machine learning process to assign daughter cells to mother cells following cell division events. Two separate modules can finally be used to visualise the output of FAST: the Overlay module plots trajectories and/or the results of analyses over the top of the original images, while the Plotting module contains a range of different options to visualise extracted data.

The FAST GUI guides the user through the process of analysing a single dataset. However, many applications require a large number of imaging datasets to be analysed using consistent settings. To automate this, we have implemented a batch-processing tool called doubleFAST. Once a user has performed an analysis on a single dataset using the FAST GUI, doubleFAST can then read the settings used during this initial run and automatically apply them to any number of additional datasets. This allows data from multiple experiments to be processed with minimal amounts of additional user input and ensures each has been analysed using consistent settings.

Finally, we have implemented a post-processing toolbox, which contains scripts and functions to perform a number of different tasks on FAST’s output. These allow, among other things, conversion of track data to other file formats, automated detection of different genotypes, and the annotation of events such as reversals in movement direction. Users of FAST can suggest new additions to this toolbox so they can be used by the wider community.

### FAST’s tracking algorithm

One of FAST’s principal innovations is its new tracking algorithm, which automatically determines how to best combine data from a variety of different cell features to improve tracking fidelity. Although previous tracking software packages have had the option to incorporate cell characteristics other than position in their tracking routines^20,22^, FAST uses a conceptual framework based on information theory that optimises this process, increasing both the power and convenience of this approach. In this section we provide a high-level overview of our main innovations and how they impact the tracking process. For more detailed derivations and explanations, please refer to section 3.4 of the Supplementary Information.

Our approach to the tracking problem is based on measurements of the amount of information that can be used to assign links between objects in subsequent frames, a measurement we call the “trackability”. This trackability score provides an integrated measure of how accurately we can follow objects from frame to frame, and due to its grounding in information theory has certain desirable properties such as the additivity of contributions from statistically independent features^27,28^. Measuring the trackability over time therefore provides users with a tool to predict when tracking will be stable and robust, as well as flagging portions of a dataset that might yield spurious trajectories. We define trackability in the following paragraphs and then illustrate how this metric is used in practice in our first case study.

We begin by assuming that *N* different features are measured for each object. An object’s features can be expressed as a vector 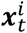 with *N* elements, where the object index is denoted as *I* and time is denoted as *t*. Changes in an objects’ features over time – corresponding to, for example, translational movement or changes in fluorescence intensity – therefore generate a trajectory through the *N-* dimensional feature space. The goal of our tracking algorithm is to reconstruct each object’s trajectory from the cloud of individual data points resulting from the segmentation and measurement of objects in each image. As previously noted, individuals in high-density and high-motility systems can easily move further than the typical cell-cell separation between frames, making it difficult to reconstruct their trajectories from positional information alone. By including additional features in the tracking framework, we expand the feature space from these two spatial dimensions to *N* feature dimensions, creating new axes along which one can potentially discriminate neighbouring individuals from each other.

The trackability metric provides an estimate of how distinguishable trajectories are from each other in the feature space, and therefore how accurate tracking is likely to be. Unpredictable movement of objects tends to reduce trackability, while trackability increases if the features sample a wider range of values (*i*.*e*. if they have a larger dynamic range). To formalise this, we model an object’s instantaneous position in feature space as the random vector ***X***_*t*_ and the change in this position between subsequent images as the random vector Δ***X***_*t*_. The distribution *f*(***x***) is then the probability density function (PDF) representing the chance of finding a randomly selected object at a particular position ***x*** in the absence of additional information (specifically, the position of the object at prior timepoints), while the distribution *f*(Δ***x*)** represents the stochastic change in an object’s position in feature space between sequential timepoints. We estimate *f*(Δ***x*)** by assuming that the motion of an object through feature space can be modelled as a Gaussian random walk. Explicitly, we assume that the feature vector 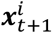 of an object at frame *t*+1 can be written in terms of its prior feature vector 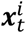 as:

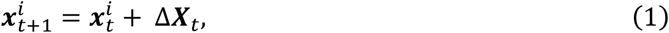

where Δ***X***_*t*_ is modelled as a multivariate normal *𝒩*(*µ*_*t*_(Δ***x***), Σ_*t*_(Δ***x***)), and *µ*_*t*_(Δ***x***) and Σ_*t*_(Δ***x***) are respectively the mean vector and covariance matrix of the set of frame-frame feature displacements {Δ***x***_*t*_}. We can similarly characterise *f*(***x***) using the covariance matrix of the raw object locations, Σ_*t*_(***x***). While we can estimate Σ_*t*_(***x***) directly from static snapshots as the covariance of the set of feature vectors {***x***_*t*_}, resolving {Δ***x***_*t*_} and subsequently *µ*_*t*_(Δ***x***) and Σ_*t*_(Δ***x***) requires a putative set of cell trajectories, which in our algorithm are obtained via a preliminary round of tracking that uses a simple nearest-neighbour approach. These statistics are robust to small numbers of tracking errors, allowing us to train our model using this relatively low-fidelity tracking approach.

From these measurements, we estimate the amount of information available for assigning objects between frames by calculating the difference between the entropies for the two distributions, H(***X***) and H(Δ***X***) ^28,29^.The trackability *r*_*t*_ in bits object^-1^ is then given as:

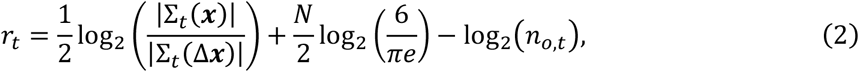

where *n*_*o,t*_ is the total number of objects present at time *t* and |·| denotes the determinant of the contents.

To illustrate the trackability metric, we consider the case of a single object with a single feature, for example its position in space along a single axis, *x* (Fig. 2a-c). This simplifies Eq. (2) to:

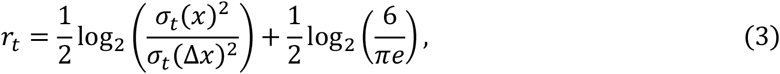

where *σ*_*t*_(*x*) and *σ*_*t*_(Δ*x*) denote the standard deviation of their respective distributions. As expected, one observes a larger trackability score when the distribution of feature displacements, *f*(Δ*x*), is more sharply peaked relative to the distribution of *f*(*x*) (Fig. 2b,c). *i*.*e*. when the typical distance an object moves between frames is small compared to the range of values of *x*. If the trackability score is sufficiently large, the risk that two objects move close to one another and become difficult to distinguish is small. Intuitively, the precision of tracking will increase as (*i*) the size of the random fluctuations in feature space decreases, (*ii*) the number of objects within a frame decreases, and (*iii*) the total size of the feature space the objects occupy (*i*.*e*. their dynamic ranges) increases. By taking these multiple factors into account, *r*_*t*_ represents an integrated measurement of the risk that a given object will be incorrectly linked to a different object in a subsequent frame.

**Fig. 2.**
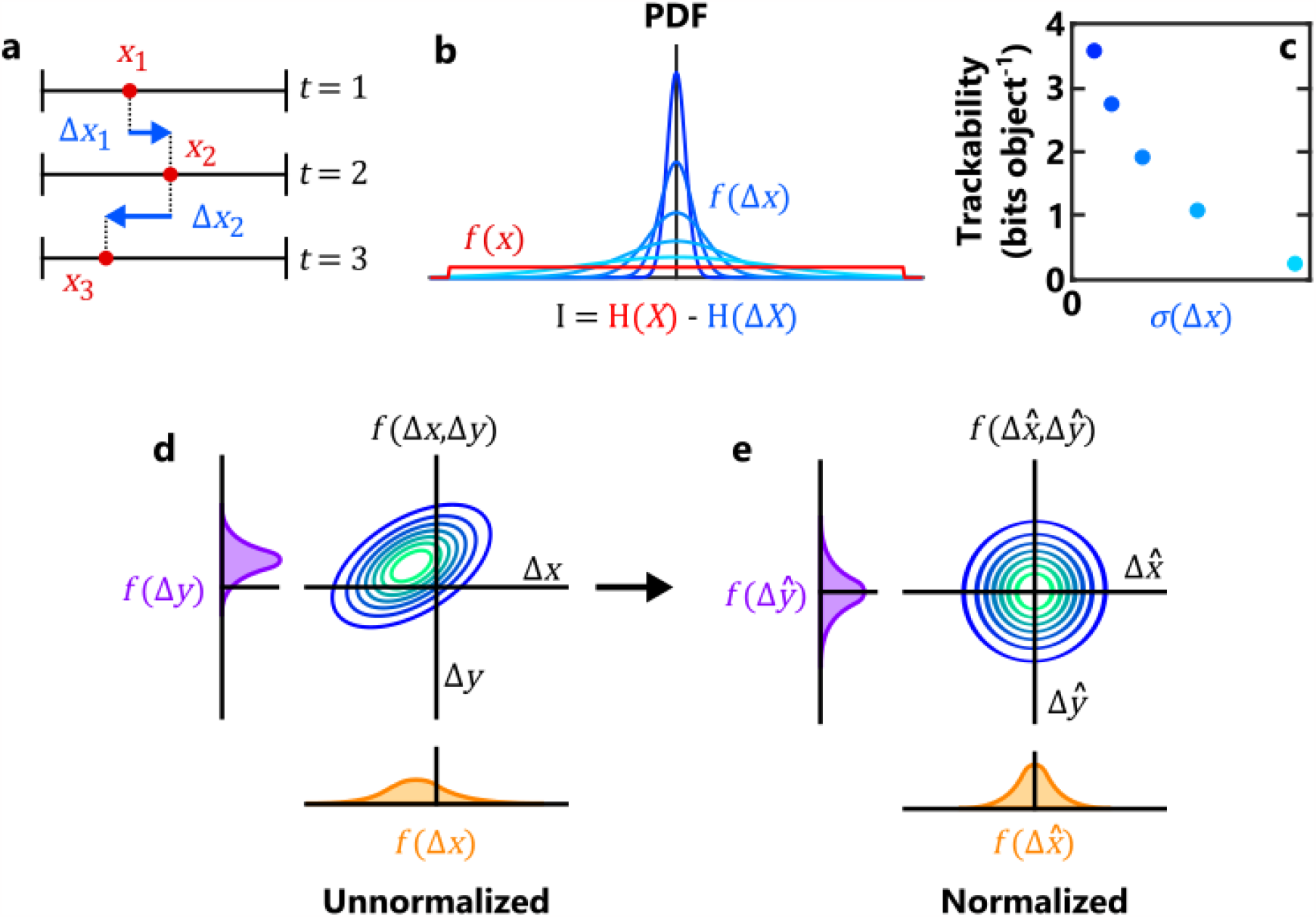
An information-theoretic framework for automated object tracking. (a) For illustrative purposes, we consider here a theoretical dataset in which an object is characterised using a single feature, its position along the x-axis. The object’s position at three successive timepoints is denoted x_1,_x_2,_x_3_ (red circles), while the displacements are denoted Δx_1,_Δx_2_ (blue arrows). We assume that feature displacements are drawn from a Normal distribution f(Δx), while the instantaneous object position (independent of knowledge of other timepoints) is drawn from a separate Uniform distribution f(x) (b). The information content I of the feature is then calculated as the difference in the entropies of the two distributions, H(X) and H(ΔX), and represents our increase in certainty about the position of the object at time t+1given knowledge about its position at time t. The trackability quantifies the total amount of information measured for each object, which increases when f(Δx) exhibits less variability relative to f(x). (c) Trackability decreases when the distribution of f(Δx) is broader (i.e. the feature becomes more “noisy”). Here the different colours correspond to the different distributions of f(Δx) shown in panel b. d,e) Illustration of the feature normalization process for two features. In both d and e, the central plot indicates the joint distribution of a pair of feature displacements, while the left and bottom plots indicate the corresponding marginal distributions. In (d), the random variables representing the unnormalized frame-frame displacements of the two features - ΔX and ΔY - are correlated and displaced from the origin. Using the joint distribution’s covariance matrix Σ(Δ**x**and mean vector µ(Δ**x**), the feature space is transformed such that the resulting joint distribution of feature displacements 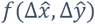 is zero-centred and isotropic (e), ensuring that each feature exhibits an equivalent amount of stochastic variation between frames.

In addition to calculating the trackability, ***µ***_*t*_(Δ***x***) and Σ_*t*_(Δ***x***) are also used to perform what we call “feature normalization**”**. This transformation converts the raw feature space ***x*** to a normalized feature space 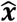 with a corresponding displacement distribution 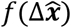 that is isotropic and zero-centred (Fig. 2d), thus ensuring that the stochastic variation observed within each component of 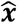 is equal and that any predictable motion in the feature space (e.g. a gradual increase in cell length due to growth, or a reduction in fluorescent intensity due to photobleaching) is accounted for. The metric of this transformed space is the Mahalanobis distance, a dimensionless measure of how reliably we can predict where an object will appear in the feature space at the next time point under the assumptions of our statistical framework. By providing an equitable way to combine data from different features together, this metric allows us to more accurately distinguish correct from incorrect putative links. Large distances between sequential timepoints in this space indicate a discrepancy between the predicted and observed location of an object in the next frame, and so suggest that the input data must be erroneous - for example, because an object was mis-segmented a single frame. In contrast to previous approaches where feature weightings have to be selected or measured manually^20^, feature normalization allows one to automatically optimise the contribution of multiple features without any additional user input.

Finally, we use these statistical measurements to also calculate the adaptive tracking threshold, which automatically adjusts the stringency of the algorithm that links objects together based on how much information is available in each frame. While a larger fraction of putative links are accepted when information is relatively plentiful, only the highest certainty links are accepted when information is more limited. This dynamic adjustment of the link threshold thus allows FAST to maximise the number trajectories in less challenging tracking conditions (*e*.*g*. slow-moving cells at low density) while minimizing the number of spurious links in more challenging conditions (*e*.*g*. fast-moving cells at high density).

### Validating the methodology of FAST using ground-truthed datasets

To demonstrate the functionality of our tracking algorithm and to illustrate how using multiple features can enhance accuracy, we used a previously described self-propelled rod (SPR) model ^9,30^ to generate synthetic data sets that simulate bacteria collectively moving at high-density (Fig. 3a). In these simulations, cells are modelled as stiff, mutually repulsive rods that are propelled by a constant force (Methods). This model has been shown to closely approximate bacterial collectives that propel themselves using either flagella or type IV pili, where steric interactions between neighbouring cells generate complex emergent collective behaviours^9,30^. Importantly, this approach allows us to independently control each of the different properties of the system, allowing us to test FAST on a large number of qualitatively different datasets and to compare the results with a known ground truth.

**Fig. 3.**
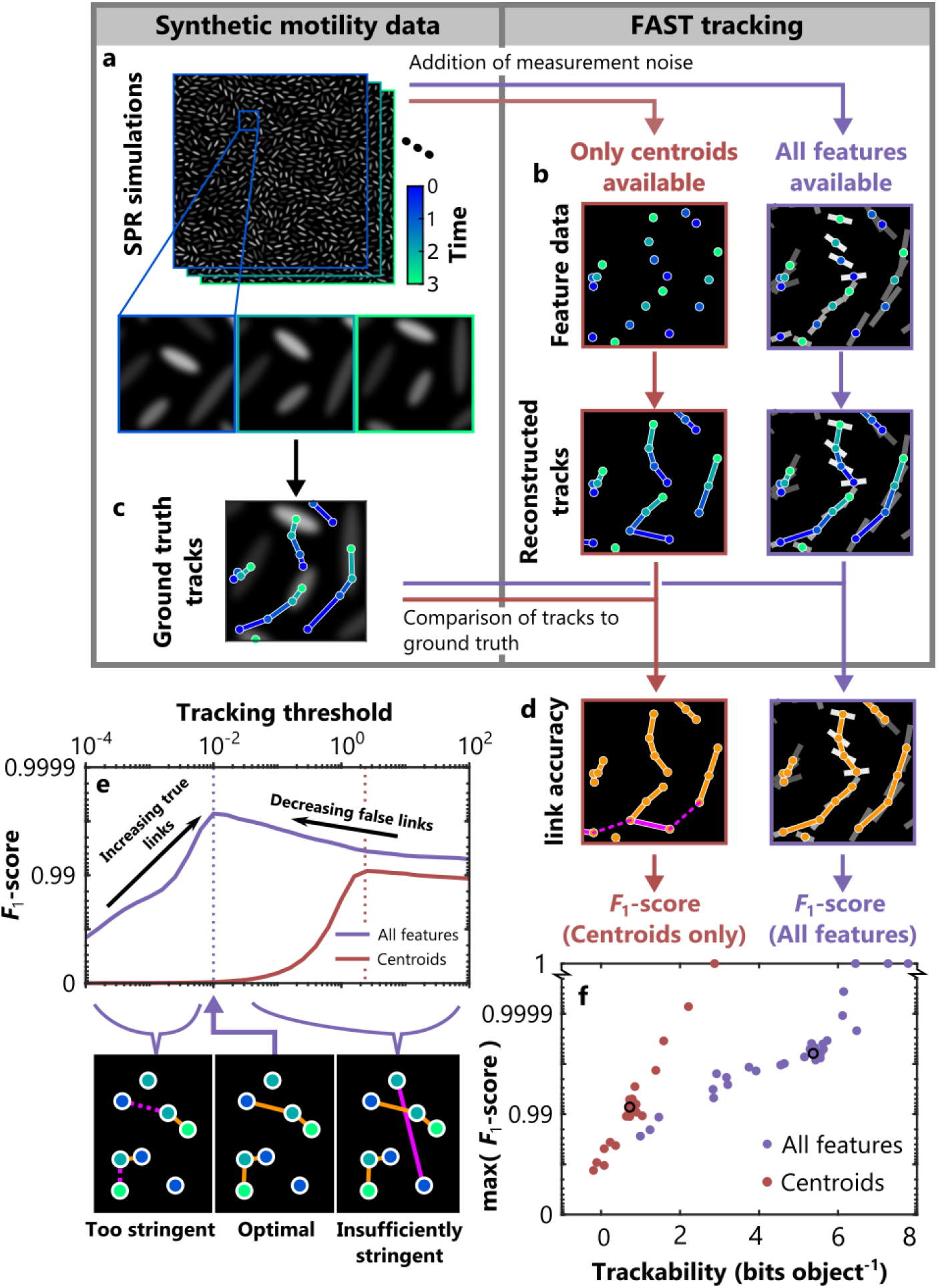
Validating FAST using synthetic data from a simulation of collective bacterial motility. (a) We used a self-propelled rod (SPR) model to generate noisy datasets with a verified ground truth. We then performed cell tracking with FAST using either only a single feature (the rod’s centroid at sequential timepoints; brown) or using a suite of different features (rod centroid, length, orientation and simulated fluorescence intensity; purple). Comparison of the reconstructed trajectories from FAST (b) with the ground truth data (c) allowed us to identify the errors made by the tracking algorithm (d). Errors were split into two categories: links made between objects that were incorrect (‘false positives’, solid magenta lines), and links between objects that were missed (‘false negatives’, dashed magenta lines). We also calculated the number of correct links made in each case (‘true positives’, solid orange lines). From these counts, we evaluated the performance of the tracking algorithm by calculating the F_1_-score (main text). (e) The value of this F_1_-score depends on a user-defined parameter, the tracking threshold. To objectively compare the results from the tracking algorithm when run on datasets with different properties, we calculated the tracking threshold that generated the largest F_1_-score for a given dataset and used this score in subsequent analyses. (f) Including all feature information substantially and consistently improved tracking performance compared to when only object positions were used (see also Extended Data Fig. 2). Furthermore, we found that our trackability metric was an excellent predictor of tracking accuracy for a given set of features, suggesting that it can be used to estimate the accuracy of the algorithm even when a ground truth is not available. In the different simulations we varied rod density, propulsive force, and framerate, as well the amount of noise in the measurement of position, length and fluorescence (Extended Data Fig. 3). The black circles in (f) correspond to the dataset presented in (e).

We simulated a 2D monolayer of cells constitutively expressing a fluorescent protein by initialising populations of cells whose length and fluorescence intensity were drawn from distributions obtained from experimental data (Extended Data Fig. 1). Following initialization, we simulated cell movement by numerically integrating the equations of motion for each rod. Once the system had reached steady-state, we extracted measurements of rod position, orientation, fluorescence and length at evenly spaced timepoints. To simulate noisy measurements, we added Gaussian noise to each of these features, with the noise magnitude based on that observed in real experimental data (Fig. 3b, Methods).

Rather than specialising on datasets with specific properties, FAST’s tracking algorithm is designed to be robust to a wide diversity of different conditions by automatically compensating via feature normalization. To test this capability, we varied the parameters of our simulation by adjusting the rod density, self-propulsion force and framerate, as well as the accuracy of feature measurement by adjusting the amount of noise in the measured cell position, length and fluorescence intensity. We tested five different values for each of these six parameters, yielding a total of 30 datasets in total.

The performance of the tracking algorithm was assessed by comparing its output to the ground truth (Fig. 3c,d). Links that were present in the ground truth but missing in the reconstructions were scored as false negatives (FN), while those that were absent in the ground truth but present in the reconstructions were scored as false positives (FP). Links that were identical in both were scored as true positives (TP). We now integrated these measurements into a single metric that quantifies tracking performance, the *F*_1_-score, defined as:

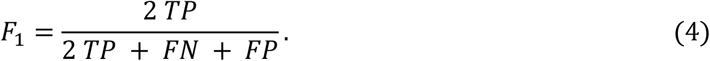

Users of FAST specify a tracking threshold that controls the stringency of the linking process. If this threshold is too stringent, too many correct links will rejected by the algorithm, while if the threshold is not stringent enough too many incorrect links will be accepted. In practice, users need the ability to choose the tracking threshold that best suits their needs - for example, users interested in detecting very rare events will have different requirements than those interested in measuring average cell behaviour. However, for the purpose of testing the benefits of our automated tracking algorithm, we removed the tracking threshold as a factor from our analyses by using the threshold that resulted in the largest the *F*_1_-score for each dataset (Fig. 3e). This greatest *F*_1_-score is an objective measure of the performance of our tracking algorithm across datasets with different characteristics, and also allows us to robustly benchmark our tracking algorithm.

The simulated datasets were tracked using two different methods (Fig. 3b). The first method only used the position of the rods – effectively a classical position-based tracking approach (‘Centroids’), while the second method used all four features – position, orientation, length and fluorescence intensity (‘All features’). We found that including all features led to a dramatic increase in tracking accuracy in all datasets, with a 4 to 10-fold reduction in the number of tracking errors (Extended Data Fig. 2). Furthermore, our groundtruthed datasets allow us to directly relate a data set’s trackability to the accuracy of the resulting tracks, showing that it is an excellent predictor of the *F*_1_-score regardless of the simulation parameters or level of noise in the dataset (Fig. 3f).

Taken together, these analyses validate our methodology and illustrate how FAST copes with challenging data sets. First, they demonstrate that our method of integrating additional feature information during tracking is reliable and effective, allowing FAST to substantially reduce tracking errors compared to alternative approaches that use cell position alone. Second, these analyses demonstrate that our trackability score is able to integrate multiple aspects of the dataset into a single, robust heuristic of predicted tracking accuracy. Finally, we note the tracking algorithm used by FAST is fully automated and requires the user to choose only a single parameter (the tracking threshold) no matter the number of different features that are used in the tracking algorithm.

### Case studies

### Quantifying rapid bursts of cell movement in densely packed *P. aeruginosa* monolayers

Many different species of bacteria generate collective motility in densely-packed communities using either flagella, Type IV pili, or via gliding^10,19,30,31^. Motility allows populations to rapidly expand into new territory, giving them a competitive advantage over non-motile genotypes^9,32^. Here we demonstrate how FAST can be used to quantify the behaviour of *P. aeruginosa* cells in interstitial colonies that form between agar and glass^31^. We focus on cells within the monolayer that forms directly behind the colony’s leading edge, the dynamics of which play a crucial role in the competition between genotypes in both interstitial and more classical “surficial” colonies^9^.

Tracking cells undergoing collective movement requires a much larger acquisition frame rate compared to non-motile cells. While expensive timing boards can be used to “trigger” cameras with a high level of precision to keep the time between frames nearly constant, most high-end research microscopes lack this capability. Instead, “camera streaming” is more widely available, which directly streams the camera’s output to a computer which saves frames as fast as possible. This maximises the frame rate, but images acquired via camera streaming can have slight variations in the time that elapses between subsequent frames.

To illustrate how FAST can be used to handle a sequence of images collected via camera streaming, we collected a large dataset with 3,505 frames recorded at an average rate of 127 frames per second. The size is of this dataset is approximately 8 Gb and each frame contains approximately 1,700 tightly packed *P. aeruginosa* cells. Despite the large size of this dataset, processing with FAST could be completed on a standard laptop computer (Methods), requiring only 200 mins to segment and 355 mins to extract the features of each of the ∼6 million individual objects.

Tracking consists of two separate stages: the model training stage, and the link assignment stage. After completion of the model training stage, FAST automatically generates and plots the trackability of the dataset at each timepoint (Fig. 4a). For our dataset, this plot revealed that the trackability dropped precipitously at some timepoints. We hypothesised that these decreases resulted from a reduction in the imaging framerate. To test this, we plotted the trackability score against Δ*t*, the elapsed time between frames as calculated from timestamps (Fig. 4a, inset). This revealed a strong negative correlation (Pearson’s correlation coefficient = -0.705), suggesting that the longer the time between frames, the lower the trackability and the less accurate the tracking results. To avoid these timepoints - and the spurious tracking results they might generate - we used the tracking module’s built-in time window selection tool to specify a subset of frames to track (Fig. 4a, green region, 750 frames). Training the tracking model for this reduced dataset took 18 minutes, while tracking and track processing took an additional 88 minutes.

**Fig. 4.**
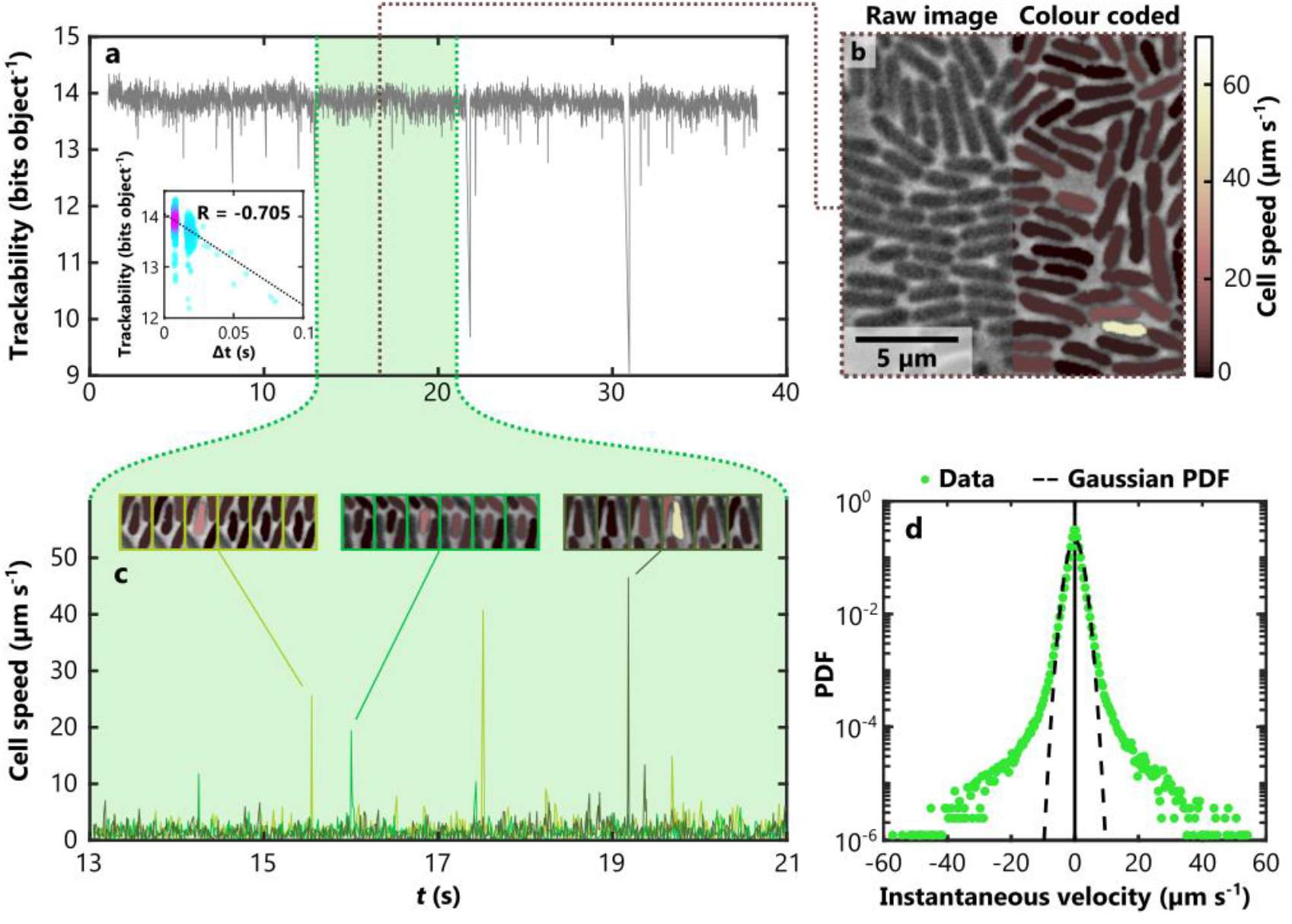
Tracking single cells in an interstitial P. aeruginosa colony spreading via pili-based motility. (a) We used FAST to track cells within a monolayer of P. aeruginosa undergoing collective motility using position, length and width as features. Images were collected using high speed “camera streaming” with a mean frame rate of 127 fps, but variations in the elapsed time between subsequent frames, Δt, resulted in transient reductions in trackability. (a, inset) Analysis of the timestamps associated with each frame revealed that the trackability score was negatively correlated with Δt (R indicates Pearson’s correlation coefficient; the warmer colours denote a higher density of data points). We therefore restricted our subsequent analyses to a subset of the data in which the trackability score was relatively constant (green region) using FAST’s built-in tools. (b) We used the Overlays module to colour code cells based on their instantaneous speed. Although cells typically moved relatively slowly, very occasionally cells were observed to undergo a very rapid burst of movement (see cream coloured cell). (c) These rapid jumps can also be observed in traces of the speed of individual cells. Here we plot the instantaneous speed of three different cells over time, each in a different colour. The montages above illustrate three of these transient events, using the same colour coding shown in b. (d) To investigate these movements at the population level, we calculated instantaneous cell velocities in both the x and y direction for all cells and plotted their combined distribution. In other active systems, this distribution is approximately Gaussian. However, in our system the highly transient bursts of velocity results in heavy tails, causing them deviate from a Gaussian distribution with the same variance (dashed black line).

In this example, a relatively large timestep Δ*t* allows cells to move further between frames, which reduces predictability and therefore reduces the trackability score. More generally, however, the trackability depends on a combination of experimental and imaging conditions, providing a robust metric to interpret datasets. For example, the trackability score can be used to quickly identify a wide range of problems which might arise during an experiment, including fluctuations in focus, illumination intensity, or inadvertent movement of the sample. Once problematic timepoints have been identified, the user can decide either to avoid them by using a subset of the images (as in this example) or to reject the entire dataset.

Tracking the smaller subset of images that we specified resulted in over 1,600 separate trajectories, each at least 200 timepoints long. To visualise this large dataset, we used the Overlays module to colour individual cells in the phase-contrast image based on their instantaneous speed (Fig. 4b). This revealed that while most cells move at relatively slow speeds, a small number of cells undergo rapid, sporadic bursts of movement approximately 25 times faster than the average. These rapid movements are also clearly visible in measurements of the speed of individual cells over time (Fig. 4c). Our results are similar to the ‘slingshots’ previously observed in solitary *P. aeruginosa* cells moving at the glass/liquid interface, apparently driven by pilus detachment events^33^. However, the peak speeds that we record (up to 60 µm s^-1^) in collectives of *P. aeruginosa* are approximately 20 times larger than those previously observed. While we do not know the specific reason for this difference, interactions with neighbouring cells facilitated by the high cell density, the glass/agar interstitial colony environment and our higher framerate (∼13 times that of ^33^) may each play a role.

To illustrate how this large tracking dataset can be mined to elucidate the statistics of rare events, we constructed the instantaneous marginal velocity distribution from our tracks (*i*.*e*. the x- and y-components of the instantaneous velocity vectors) (Fig. 4d). While previous studies have found that sperm and swimming bacterial cells undergoing collective movement generate marginal velocity distributions that are approximately Gaussian^30,34^, we observed that bacteria collectively moving via pili-based motility exhibit much heavier tails corresponding to their occasional rapid motions. Despite the rarity of these events – less than 0.05% of our measurements had a magnitude larger than 20 µm s^-1^ - the exceptional number of cell trajectories we obtained nevertheless allowed us to finely resolve their statistical distribution. These analyses demonstrate how FAST can be used to rapidly characterise the motility of a large number of cells in densely-packed conditions, with relatively little user input and computational effort.

### Lineage analysis of *Escherichia coli*

A major aim of a number previous software packages has been to automatically reconstruct cell lineages^5,8^. However, identifying cell division events and the resulting daughter cells is a particularly difficult challenge. Near-perfect tracking of individuals is essential for accurate lineage tracking, as a single error can propagate through multiple division events, often contaminating the remainder of the dataset. We decided to test FAST’s lineage tracking capabilities using a time lapse movie of dividing *E. coli* cells^17^ imaged using phase-contrast microscopy. Although the tracking algorithm performed well, several links were incorrectly assigned – this was particularly problematic at the beginning of the dataset, because the small number of cells provided insufficient data to adequately train the machine learning model. We therefore performed manual correction of the dataset using FAST’s built-in track validation and correction system, which required us to break 21 incorrect links and form 17 new links. These manual corrections constituted only 0.67% of the 5,698 links in the final dataset, with the remainder of links being correctly assigned automatically.

Following automated division detection, we visualised the lineage structure of the colony using the Overlays module (Fig. 5a, b, Supplementary Video 1), allowing us to rapidly verify the accuracy of the lineage assignment. Visualisation of the population structure revealed that the descendants of each of the individual cells present at the beginning of the experiment formed highly elongated structures within the colony, similar to the patterns previously observed in experiments where colonies were initiated from multiple founder cells labelled with different fluorescent proteins^14^. We also used the feature extraction and plotting modules to reconstruct the length of individual cells and align them by the time that they last underwent division (Fig. 5c). These analyses illustrate the power and accuracy of our tracking and lineage reconstruction approaches, and also showcase the ease with which FAST allows data-rich visualisations of the resulting datasets.

**Fig. 5.**
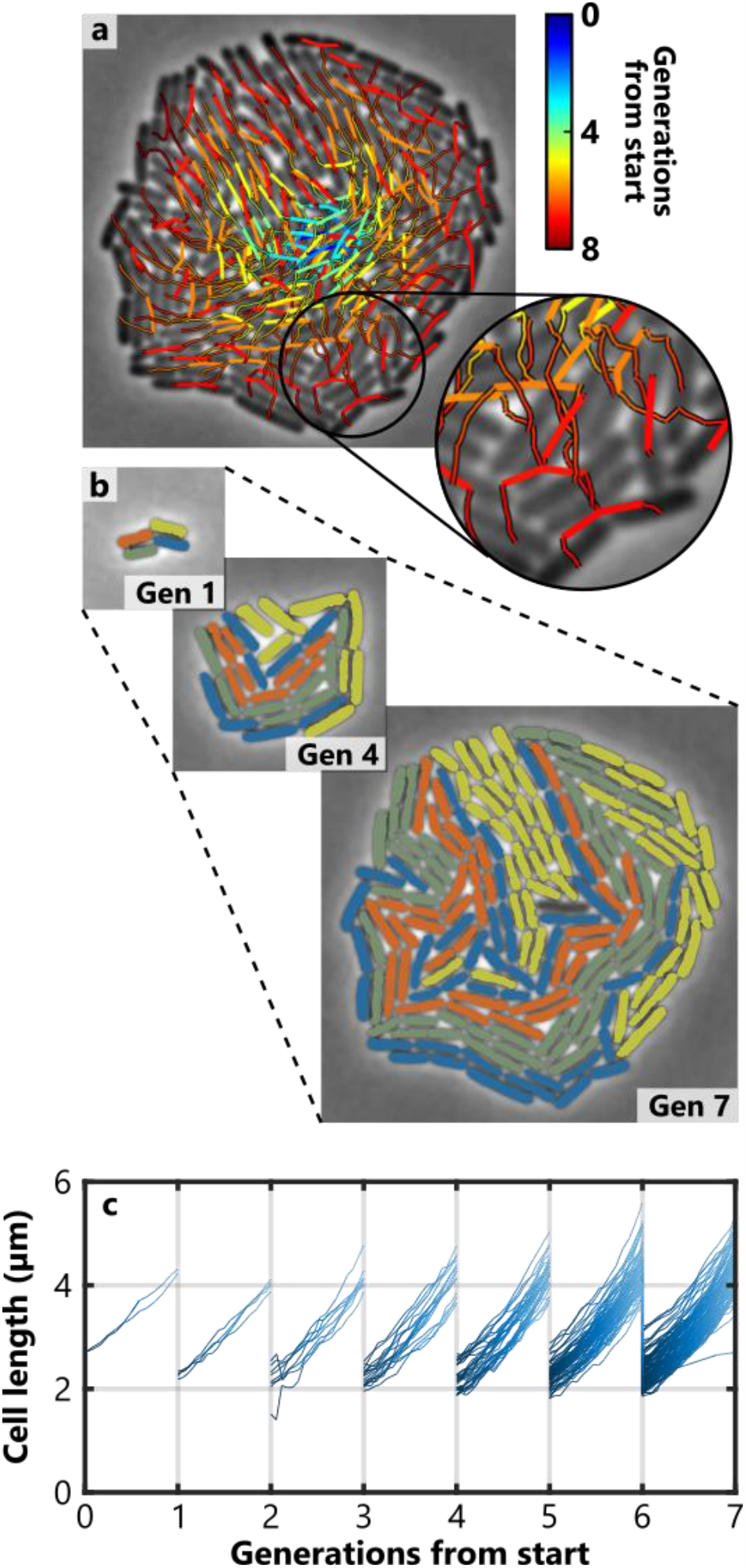
Tracking and lineage analysis of E. coli cells in a growing colony. (a) Lineage tree showing the colony at the final point of the timeseries. Here thicker lines show the location of division events and thinner lines show the movement of cells between divisions. The trajectory colour indicates how many generations have elapsed in each lineage from the initial timepoint. (b) The colony at three different timepoints. Cells that share the same mother in generation 1 (‘Gen 1’) are shown in the same colour at each timepoint, illustrating how the spatial distribution of different lineages develops over time. (c) Growth curves of each cell, aligned by generation number. Tracks are shaded according to the age of the corresponding cell, from birth (dark blue) to division (light blue). The data visualisations of both panels a and b were produced using FAST’s Overlays module, while panel c was prepared with the Plotting module.

### Automated analysis of T6SS battles

To illustrate FAST’s capabilities beyond standard cell tracking, we analysed the activity of the Type 6 Secretion System (T6SS) using FAST. The T6SS is a highly dynamic organelle composed of a molecular ‘spear’ tipped with a toxin known as an effector^35^. Bacteria that possess the T6SS can inject this effector into neighbouring cells, which kills them^36^. Firing events can be monitored by visualising the localisation of the protein that forms the contractile sheath of the T6SS needle, TssB in *P. aeruginosa* and VipA in *Vibrio cholerae*^37,38^. Here, we highlight how FAST can be used to quantify how the T6SS is regulated in co-cultures of *V. cholerae* and *P. aeruginosa*.

Because of its short range, the T6SS is effective only at high cell density, which historically has made it difficult to study using automated analyses. We used FAST to analyse imaging datasets that show two different bacterial species interacting with their respective T6SS when mixed together in a densely packed monolayer^37^, specifically *V. cholerae* and *P. aeruginosa* cells expressing VipA-mCherry and TssB-mNeongreen, respectively (Fig. 6a, left). We distinguished the two species from one another (Fig. 6a, right) using FAST’s post-processing toolbox, which compares each cell’s intensity in the mCherry and mNeonGreen channels to automatically assign them to two distinct populations (Fig. 6b). This utility allows one to rapidly compare the behaviour of different genotypes within the same sample.

**Fig. 6.**
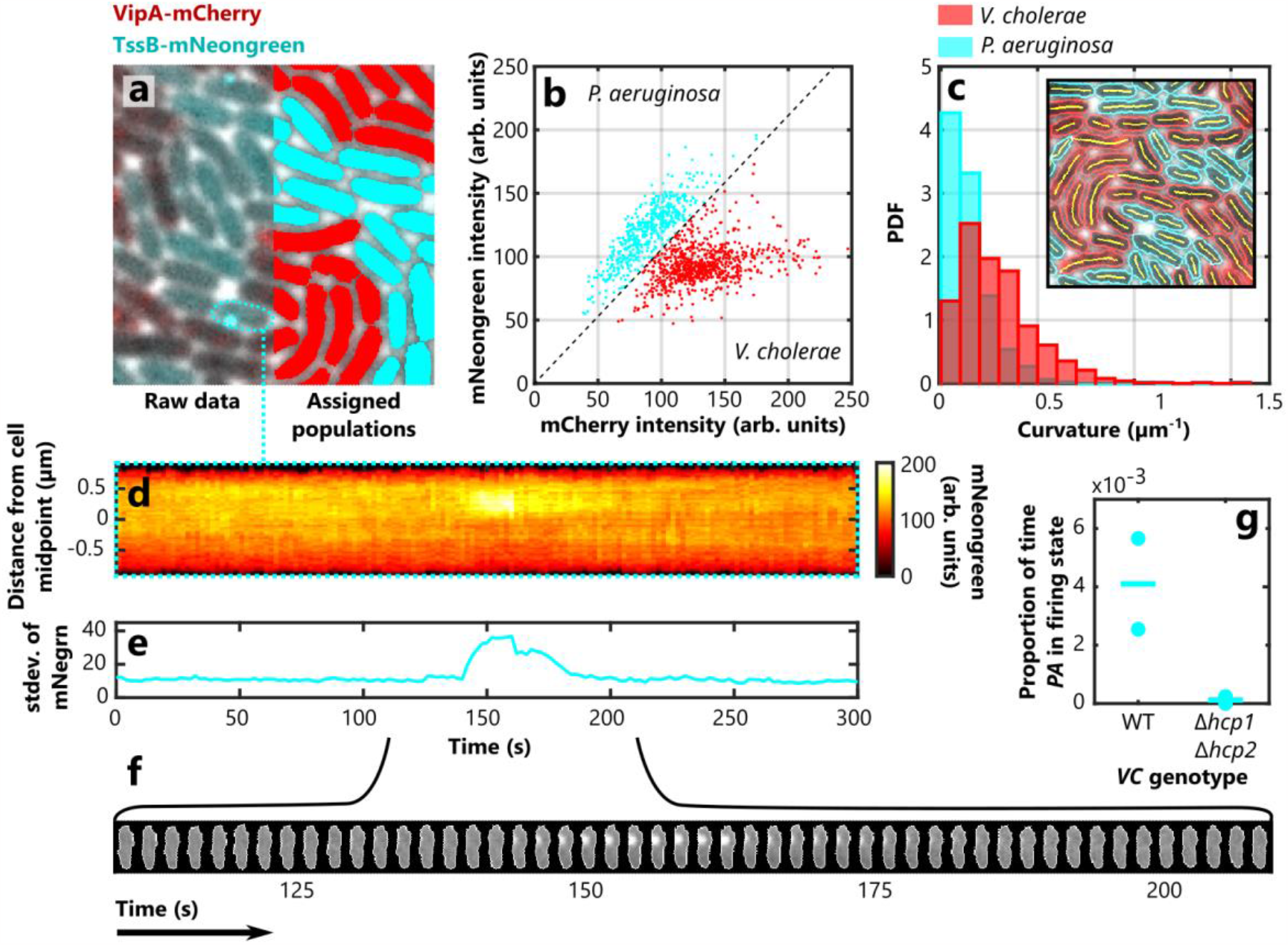
Identifying different bacterial species, quantifying cell morphologies, and investigating T6SS dynamics in a densely packed collective. (a) A monolayer composed of a mixture of P. aeruginosa cells expressing TssB-mNeongreen and V. cholerae cells expressing VipA-mCherry (left). Measurements of the average intensity of each cell in the two different fluorescence channels allowed us to automatically distinguish the two species (b), an approach known as image cytometry. By colour coding each cell (P. aeruginosa, cyan, V. cholerae, red), we were able to visually confirm the accuracy of the assignment process (a, right). (c) We then quantified a novel feature, cell curvature, using FAST’s custom feature extraction framework to calculate the curvature of the segmentation backbones (inset). Splitting these measurements into the two previously assigned populations showed the comma shaped V. cholerae cells were substantially more curved than the rod-shaped P. aeruginosa cells. (d-f) We then used FAST’s plotting options to illustrate the dynamics of the T6SS. A P. aeruginosa cell fires its T6SS machinery (observed as the bright puncta of TssB), which can be visualised using the kymograph (d) and ‘cartouche’ (f) plotting options, as well as an increase in the standard deviation of the mNeongreen channel (e). (g) We used this latter measurement to detect when the T6SS was active in a cell, allowing us to measure the proportion of time that each cell in the population spends in the firing state. Our analyses confirmed that when co-cultured with a strain of V. cholerae with an inactive T6SS (Δhcp1 Δhcp2), the T6SS activity of WT P. aeruginosa cells is dramatically reduced compared to when they are co-cultured with WT V. cholerae. Here the circles show results from individual experimental timeseries and horizontal lines show the overall mean proportion.

The architecture of FAST allows users to easily define and quantify novel features that are not already included in FAST. To do this, users can write a short feature measurement script that can make use of the segmentation of each object in each frame, as well as the corresponding regions in each of the imaging channels - in this case, the mNeongreen, mCherry and phase-contrast signals. The Feature Extraction module then automatically stores these custom features in the same way as the built-in features, allowing them to be integrated into downstream analyses. To illustrate this capability, we extracted cell curvature as a custom feature to see if we could detect the differences in morphology of *V. cholerae* cells (which are generally comma-shaped) from that of *P. aeruginosa* (which are generally rod-shaped). Using a skeletonization-based approach, we extracted the morphological backbone of each cell from its segmentation and measured its curvature by finding the best-fit circle to this set of points. Consistent with expectations, our automated analysis found that the cells identified as *V. cholerae* were substantially more curved than those identified as *P. aeruginosa* (Fig. 6c).

Our software is also capable of generating sophisticated visualisations and analyses of the behaviour of single cells. For example, FAST allows the dynamics of T6SS firing in individual cells to be easily visualised using the kymograph option of the Plotting module (Fig. 6d), as well as the ‘cartouche’ option, which extracts and aligns cropped images of the specified cells (Fig. 6f). We can also quantify the dynamics of the T6SS using the feature data associated with a given cell: the standard deviation of the mNeongreen channel substantially increases during firing (Fig. 6e), as the TssB becomes non-uniformly distributed through the cell as it assembles into the sheath.

It is generally accepted that *P. aeruginosa* only fires its T6SS when triggered by the firing of the T6SS of a neighbouring cell^15,37^. Although qualitative evidence for this is strong, to our knowledge this effect has never been directly quantified, likely because of the difficulties involved with tracking densely packed cells and because the T6SS firing events are themselves relatively rare. We therefore decided to use FAST to simultaneously track a large number of densely packed cells and automatically detect when each fires its T6SS. We used our batch-processing tool – doubleFAST – to automate the analysis of multiple datasets from experiments in which WT *P. aeruginosa* cells were either cocultured with WT *V. cholerae* or cocultured with a mutant *V. cholerae* strain that is incapable of firing their T6SS (Δ*hcp1* Δ*hcp2*). Using the standard deviation of the mNeongreen channel to distinguish when *P. aeruginosa* is actively firing its T6SS, we calculated the proportion of time that *P. aeruginosa* cells spend firing their T6SS (Fig. 6g). As expected, when co-cultured with the inactivated Δ*hcp1* Δ*hcp2 V. cholerae* strain, we measured that *P. aeruginosa* reduced the proportion of time it spent in the firing state approximately 30-fold compared to when co-cultured with WT *V. cholerae*. In total, these analyses were based on 1,246 tracks built from 139,849 individually segmented objects, illustrating how our automated processes can leverage quantities of data not amenable to manual approaches. In conclusion, this case study demonstrates how FAST can automatically distinguish different species, how users can specify novel features like cell curvature, and how complex bacterial behaviours can be quantified using feature data.

## Discussion

Advances in experimental techniques and automated microscopy have transformed live cell imaging from a technique that relies largely on qualitative observation into a highly quantitative discipline that leverages huge amounts of data. Integral to this renaissance are computational tools that can automatically parse and annotate the large imaging datasets resulting from these experiments. Many tools have been developed for this purpose^3–8,39–44^, each optimised for a specific research question. As we have emphasized throughout this manuscript, our own contribution is particularly well-suited for the analysis of experiments where cells are tightly packed together.

We designed FAST to allow users to rapidly obtain high-quality results by minimising both the number of user-defined tracking parameters and the computational time required to process an imaging dataset containing many cells. FAST automatically determines how to combine data from different cell features, which prevents users from having to iteratively improve tracking results by sequentially adjusting a large number of parameters – typically a very slow and laborious process. Our approach also provides the user with a single, easy to interpret metric that allows them to rapidly identify and avoid sections of the dataset that are predicted to yield low-fidelity trajectories. While the basic FAST package already outputs most of the cell features that are widely used by researchers, simple modifications to the system allow users to integrate new features into the existing analytical framework.

Tracking systems typically break down in one of two ways – either individual objects cannot be distinguished, or the tracking algorithm cannot accurately link individuals between frames. Machine learning is increasingly used to solve the first of these two challenges, and techniques that apply machine learning methods to segment microbes at high density are now widely available^41–43,45^. However, use of machine learning to perform tracking is comparatively rare. FAST fills this niche using a statistical framework that both substantially improves track quality and provides users with insight into possible problems with their data. Our software has already proven its ability to elucidate the rich and complex behaviours that individual bacteria exhibit in densely-packed conditions^9,16,25,26^. Future work stemming from these combined experimental and analytical approaches could ultimately shed new light on how dense bacterial communities function and help us to develop novel strategies to manipulate them.

## Materials and methods

### Computational resources

All analyses presented in this manuscript were performed with a Microsoft Surface Book 2, with an 8-core Intel i7-8650U CPU and 8 Gb of RAM. FAST was run directly in Matlab, version 2018b.

### SPR model

The 2D SPR model used in this manuscript has been described in detail elsewhere^9,30^. In brief, we model cells as stiff rods composed of evenly-spaced Yukawa segments, mutually repulsive point potentials. We initialise the system by filling a square domain with *N*_*r*_ rods, evenly spaced on a lattice. Each rod *i* is associated with an aspect ratio *a*^*i*^ and a fluorescence intensity *I*^*i*^, randomly drawn from distributions fitted to the *P. aeruginosa* data from the T6SS visualisation experiment^15^ as shown in Extended Data Fig. 1. The packing fraction of the system, *ρ*, is calculated as

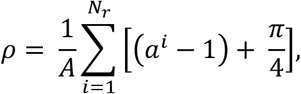

where *A* is the area of the simulated domain, which has doubly periodic boundary conditions.

Taking the instantaneous position of a rod *i* as ***r***^*i*^, its orientation as *ϕ*^*i*^, the unit vector denoting this orientation as 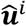 and the sum of the potentials between *i* and all other rods as *U*^*i*^, we define the equations of motion for each rod as:

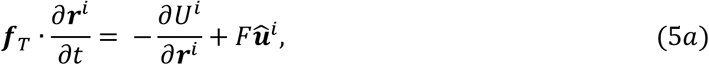

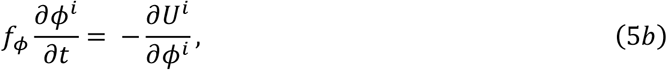

where ***f***_*T*_ is the translational friction tensor, *f*_*θ*_ is the rotational friction constant and *F* is the size of a self-propulsion force exerted by each rod along its axis. We use the formulation presented in^30^ to calculate ***f***_*T*_ and *f*_*ϕ*_, which are in turn functions of the rod aspect ratio and the Stokesian friction coefficient, *f*_*0*_. We simulate the dynamics of our system by numerically integrating Eq. 5a and 5b using the midpoint method.

Following an initial transient, the system reaches a statistical steady-state. At this time, we begin to measure the positions, orientations, lengths and intensities of each rod at a sampling “framerate” Δ*T*. We add simulated measurement noise to each of these measurements, which is drawn from a Gaussian distribution with a mean of zero. The standard deviation of the measurement noise for each feature is controlled by the parameters *σ*_*r*_ (positional noise), *σ*_*ϕ*_ (orientational noise), *σ*_*a*_ (length noise) and *σ*_*I*_ (fluorescence noise). To constrain the baseline estimates of these noise parameters, we measured how each of the corresponding features fluctuated about the mean in *P. aeruginosa* cells in a dataset of T6SS firing^15^. The cells in this dataset are non-motile and the framerate is high enough that growth is negligible, meaning any apparent changes in position, orientation, length or fluorescence are wholly attributable to measurement noise.

We varied the properties of our simulations by adjusting the values of the parameters *N, F*, Δ*T, σ*_*r*_, *σ*_*a*_ and *σ*_*I*_ in different simulation runs. The values of these parameters, as well as the fixed system parameters, are provided in table 1.

**Table 1.** Parameters of the SPR model used to generate synthetic datasets. Here we show both range of values we tested and baseline value that was used when we varied another parameter. Note that our simulations are non-dimensionalised, using the width of a single rod as the characteristic lengthscale and the time taken for an isolated rod with F=1 to move a single rod width as the characteristic timescale.

| Parameter symbol | Parameter name | Value(s) |
| --- | --- | --- |
| $A$ | Area of simulation domain | 10,000 |
| $f_0$ | Stokesian friction coefficient | 1 |
| $N$ | Number of rods | 700 (baseline)<br>[300, 500, 700, 900, 1100] (variable) |
| $F$ | Self-propulsion force | 1 (baseline)<br>[0.7, 0.85, 1, 1.15, 1.3] (variable) |
| $\Delta T$ | Time between sampled timepoints | 10 (baseline)<br>[5, 7.5, 10, 12.5, 15] (variable) |
| $\sigma_r$ | Standard deviation of positional measurement noise | 0.02 (baseline)<br>[0.02, 0.1, 0.2, 0.4, 0.6] (variable) |
| $\sigma_\phi$ | Standard deviation of orientational measurement noise | 0.02 radians |
| $\sigma_a$ | Standard deviation of length measurement noise | 0.1 (baseline)<br>[0.05, 0.1, 0.2, 0.4, 0.6] (variable) |
| $\sigma_I$ | Standard deviation of fluorescence measurement noise | 1 A.U. (baseline)<br>[0.5, 1, 2, 4, 6] A.U. (variable) |

### Sample preparation

The monolayer of *P. aeruginosa* presented in Fig. 4 was prepared using the wild-type PAO1 strain^46^. Cells were streaked out from freezer stocks onto LB agar plates (Lennox, 20 g/l, Fisher Scientific, solidified with 1.5% (w/v) agar, Difco brand, BD) and incubated overnight at 37°C. Single colonies were picked from the resulting plates and incubated overnight in liquid culture under continuous shaking, resulting in stationary phase cultures. These were then diluted 30-fold and returned to the shaking incubator for a further two hours, yielding cultures in exponential phase. The final culture used for inoculation was prepared by adjusting the optical density at 600 nm (OD_600_) of the exponential phase cultures to 0.05 using fresh LB, corresponding to an approximate concentration of 12,500 cells μl^-1^

We prepared monolayers from these cultures using a similar protocol to that described in^31^. 1 μl of inoculation culture was spotted onto the centre of a small (2 cm x 2 cm) LB agar pad. To provide optimal conditions for observing twitching motility, the concentration of agar in these pads was 0.8%. This pad was then inverted and placed into the base of a coverslip-bottomed Petri dish (175 µm coverslip thickness, MatTek), which was then closed and incubated for 16 hr at room temperature. The ability to close the lid of the Petri dish allowed us to avoid desiccation of the sample during incubation. By the end of the incubation period, a large interstitial colony with a dense monolayer at its perimeter had formed^9^.

### Microscopy

The high framerate movie used for the analysis of rapid motion (Fig. 4) was acquired using a Zeiss Axio Observer.Z1 microscope outfitted with an Axiocam 702 camera set to “camera streaming” mode and a Plan Apochromat 63x oil-immersion objective.

### Other datasets

The datasets showing dividing *E. coli* cells (Fig. 5) and T6SS firing (Fig. 6) were downloaded from the Supplementary Information sections of^17^ and^37^, respectively, under a Creative Commons Attribution license. FIJI’s^47^ built-in AVI reader was used to extract individual frames of both movies. The *E. coli* movie was then stabilised using a combination of manual stabilisation (for large shifts in the field of view between timepoints) and the TurboReg plugin^48^.

## Acknowledgements

We would like to thank the numerous people who have tested and provided feedback on FAST during its development, including N.A. Costin, S.C. Booth, J.H. Wheeler, E.T. Granato, R.K. Kumar, W.P.J. Smith, C.J. Walther and H. Todorov. O.J.M. was supported by an EPSRC studentship through the Life Sciences Interface Centre for Doctoral Training (EP/F500394/1) and a long-term postdoctoral fellowship provided by the HFSP (LT0020/2022-L), while W.M.D was supported by a startup grant from the University of Sheffield’s Imagine: Imaging Life initiative, an EPSRC Pump Priming Award (EP/M027430/1), a BBSRC New Investigator Grant (BB/R018383/1), and the Human Frontier Science Program (RGY0080/2021).

## Code availability statement

All source code, including both FAST and the SPR benchmarking system, is available at https://github.com/Pseudomoaner/FAST.

## Data availability statement

Data for the case studies, including raw data files, track data and data processing settings and scripts, are available at https://doi.org/10.6084/m9.figshare.21756212 (Fig. 4), https://doi.org/10.6084/m9.figshare.21756269 (Fig. 5) and https://doi.org/10.6084/m9.figshare.21756329 (Fig. 6)

## Conflict of interest

We report no conflicts of interest.

---

**Extended Data Fig. 1.**
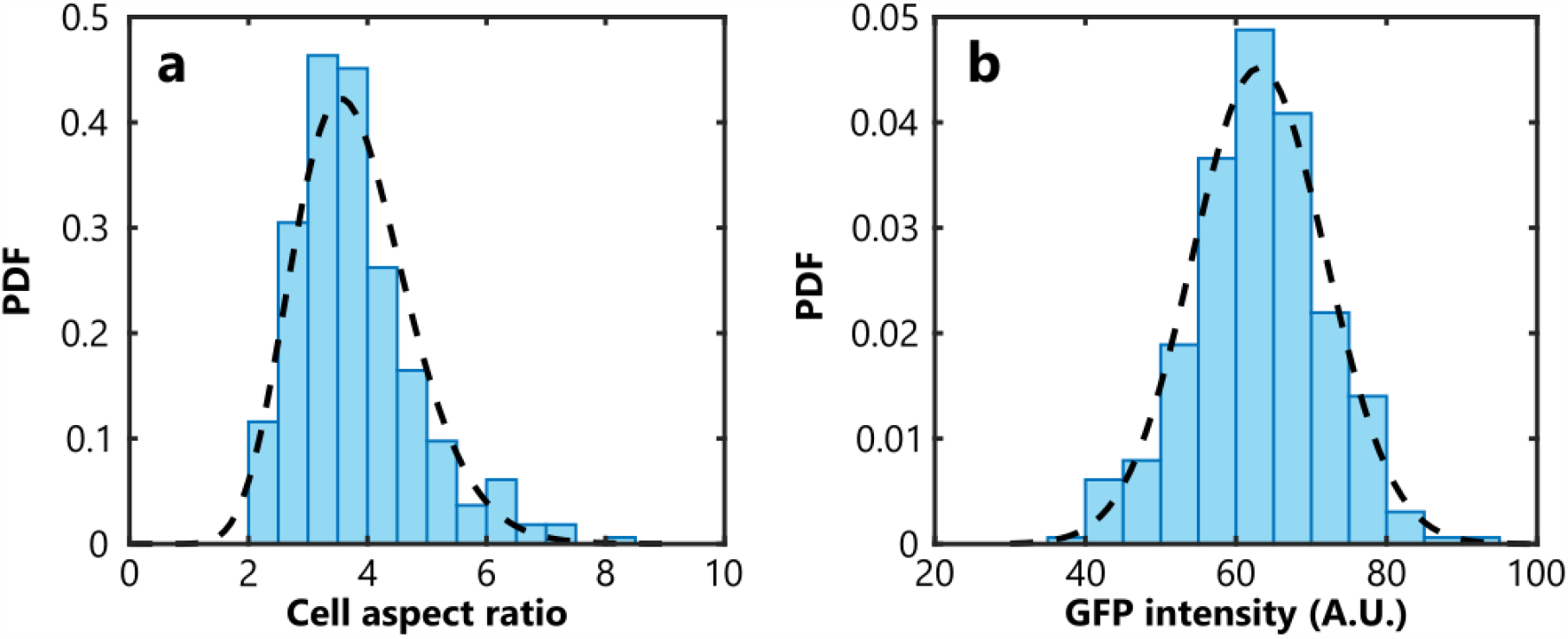
Distributions used to specify initial conditions of the SPR model. The distributions of trajectory-averaged aspect ratios (a) and GFP intensities (b) of a dataset of non-motile P. aeruginosa cells^15^. A gamma distribution (shape parameter = 15.3, scale parameter = 0.248) and a normal distribution (mean = 63.1, standard deviation = 8.83), respectively, were fitted to these two datasets (black dotted lines). To initialise the SPR model, rod aspect ratio, a_α_, and simulated fluorescence intensity, I_α_, were randomly drawn from these two fitted distributions, allowing us to ensure that these two features were modelled realistically in our simulations.

**Extended Data Fig. 2.**
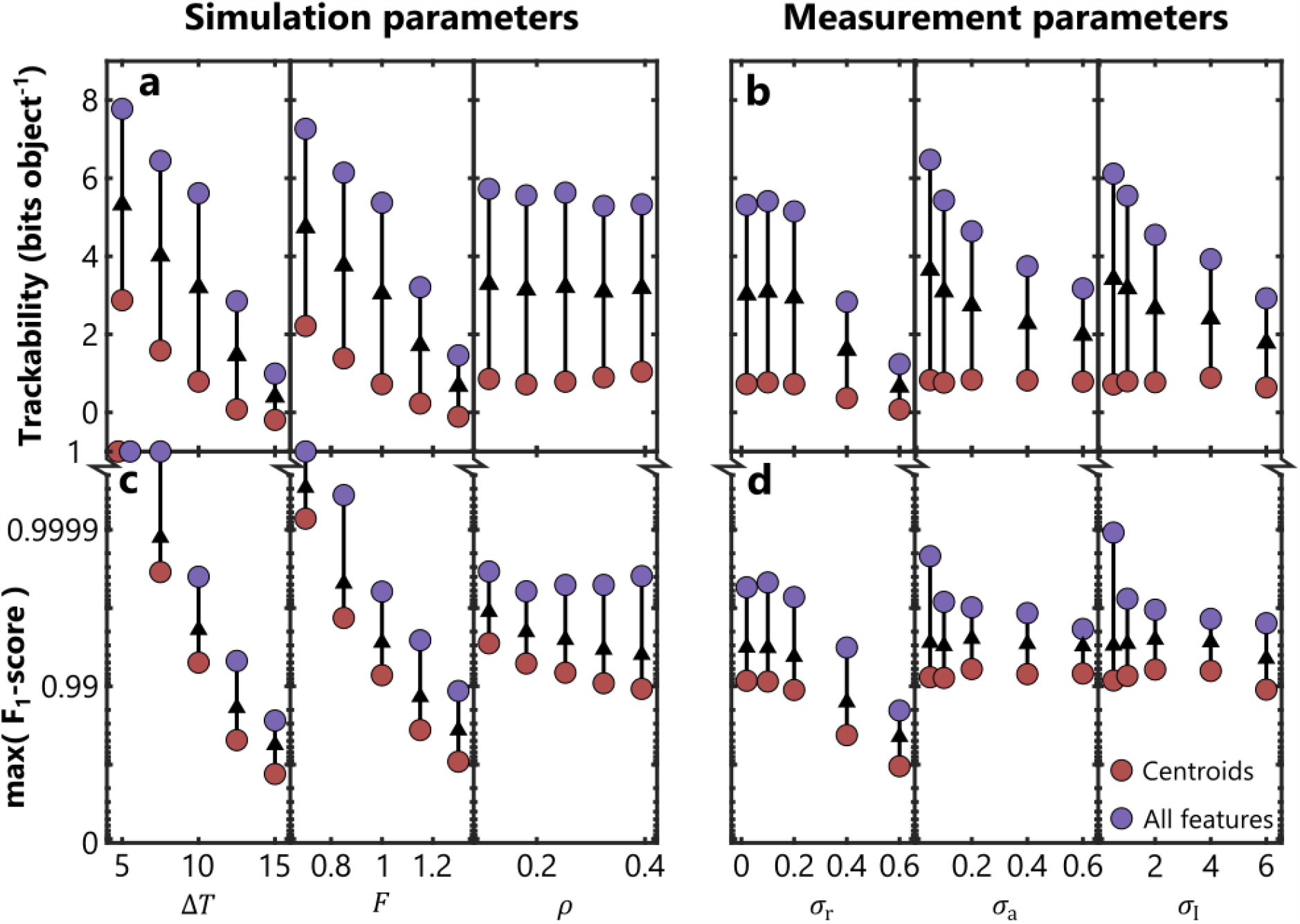
Including additional feature information improves trackability and increases tracking fidelity. We measured the trackability (a, b) and maximum F_1_-scores (c, d) of synthetic high-density motility data generated using a range of different parameter combinations. Both ‘simulation parameters’ (parameters that change the properties of the SPR model we used to generate the synthetic dataset, a, c) and ‘measurement parameters’ (parameters that change the amount of measurement noise for each of the different features, b, d) were varied. See Table 1 for further details. Here we compare trackability and tracking fidelity when the tracking algorithm can only use positional information (‘Centroids’, brown) to when all feature information is available (‘All features’, purple). Arrows show the consistent increase in these two metrics when all features from the synthetic dataset are used, illustrating the robustness of our approach.

## Notes

https://bit.ly/3vovDHn

https://figshare.com/projects/FAST_demonstration_data/155852

https://github.com/Pseudomoaner/FAST

### Competing Interest Statement

The authors have declared no competing interest.

### Summary of Updates

This new version contains a new version of figure 6 based on a different collection of datasets, as well as links to repositories where processed data presented in this manuscript can be accessed. We have added a few references to reflect recent advances in the field.

## References

1. Nadell, C. D., Xavier, J. B. & Foster, K. R. The sociobiology of biofilms. FEMS Microbiol Rev 33, 206–24 (2009).

2. Nadell, C. D., Drescher, K. & Foster, K. R. Spatial structure, cooperation and competition in biofilms. Nat Rev Microbiol 14, 589–600 (2016).

3. Ducret, A., Quardokus, E. M. & Brun, Y. v. MicrobeJ, a tool for high throughput bacterial cell detection and quantitative analysis. Nat Microbiol 1, 1–7 (2016).

4. Paintdakhi, A. et al. Oufti: an integrated software package for high-accuracy, high-throughput quantitative microscopy analysis. Mol Microbiol 99, 767–777 (2016).

5. Stylianidou, S., Brennan, C., Nissen, S. B., Kuwada, N. J. & Wiggins, P. A. SuperSegger : robust image segmentation, analysis and lineage tracking of bacterial cells. Mol Microbiol 102, 690–700 (2016).

6. Tinevez, J.-Y. et al. TrackMate: An open and extensible platform for single-particle tracking. Methods 115, 80–90 (2017).

7. Wang, Q., Niemi, J., Tan, C.-M., You, L. & West, M. Image segmentation and dynamic lineage analysis in single-cell fluorescence microscopy. Cytometry Part A 77A, 101–110 (2009).

8. Klein, J. et al. TLM-tracker: Software for cell segmentation, tracking and lineage analysis in time-lapse microscopy movies. Bioinformatics 28, 2276–2277 (2012).

9. Meacock, O. J., Doostmohammadi, A., Foster, K. R., Yeomans, J. M. & Durham, W. M. Bacteria solve the problem of crowding by moving slowly. Nat Phys 17, 205–210 (2021).

10. Jeckel, H. et al. Learning the space-time phase diagram of bacterial swarm expansion. Proc Natl Acad Sci U S A 116, 1489–1494 (2019).

11. Granato, E. T., Meiller-Legrand, T. A. & Foster, K. R. The Evolution and Ecology of Bacterial Warfare. Current Biology 29, R521–R537 (2019).

12. Smith, P. & Schuster, M. Public goods and cheating in microbes. Current Biology vol. 29 R442–R447 Preprint at https://doi.org/10.1016/j.cub.2019.03.001 (2019).

13. Madsen, J. S., Burmølle, M., Hansen, L. H. & Sørensen, S. J. The interconnection between biofilm formation and horizontal gene transfer. FEMS Immunol Med Microbiol 65, 183–195 (2012).

14. Rudge, T. J., Federici, F., Steiner, P. J., Kan, A. & Haseloff, J. Cell polarity-driven instability generates self-organized, fractal patterning of cell layers. ACS Synth Biol 2, 705–714 (2013).

15. Basler, M., Ho, B. T. & Mekalanos, J. J. Tit-for-Tat: Type VI Secretion System Counterattack during Bacterial Cell-Cell Interactions. Cell 152, 884–894 (2013).

16. Granato, E. T. & Foster, K. R. The Evolution of Mass Cell Suicide in Bacterial Warfare. Current Biology 30, 1–8 (2020).

17. Nghe, P. et al. Microfabricated Polyacrylamide Devices for the Controlled Culture of Growing Cells and Developing Organisms. PLoS One 8, e75537 (2013).

18. Kurtuldu, H., Guasto, J. S., Johnson, K. A. & Gollub, J. P. Enhancement of biomixing by swimming algal cells in two-dimensional films. Proc Natl Acad Sci U S A 108, 10391–5 (2011).

19. Jarrell, K. F. & McBride, M. J. The surprisingly diverse ways that prokaryotes move. Nat Rev Microbiol 6, 466–476 (2008).

20. Al-Kofahi, O. et al. Automated Cell Lineage Construction: A Rapid Method to Analyze Clonal Development Established with Murine Neural Progenitor Cells. Cell Cycle 5, 327–335 (2006).

21. Veenman, C. J., Reinders, M. J. T. & Backer, E. Resolving motion correspondence for densely moving points. IEEE Trans Pattern Anal Mach Intell 23, 54–72 (2001).

22. Meijeringa, E., Dzyubachyka, O., Smala, I. & Cappellen, W. A. van. Tracking in cell and developmental biology. Semin Cell Dev Biol 20, 894–902 (2009).

23. Zhao, K. et al. Psl trails guide exploration and microcolony formation in Pseudomonas aeruginosa biofilms. Nature 497, 388–91 (2013).

24. Luo, Y. et al. A hierarchical cascade of second messengers regulates Pseudomonas aeruginosa surface behaviors. mBio 6, e02456–14 (2015).

25. Krishna Kumar, R. et al. Droplet printing reveals the importance of micron-scale structure for bacterial ecology. Nat Commun 12, 1–12 (2021).

26. Ghosh, D. & Cheng, X. To cross or not to cross: Collective swimming of Escherichia coli under two-dimensional confinement. Phys Rev Res 4, 023105 (2022).

27. Dubuis, J. O., Tkacik, G., Wieschaus, E. F., Gregor, T. & Bialek, W. Positional information, in bits. Proc Natl Acad Sci U S A 110, 16301–16308 (2013).

28. Cover, T. M. & Thomas, J. A. Elements of Information Theory. Elements of Information Theory (John Wiley & Sons, Inc., 1991). doi:10.1002/0471200611.

29. Ahmed, N. A. & Gokhale, D. v. Entropy Expressions and Their Estimators for Multivariate Distributions. IEEE Trans Inf Theory 35, 688–692 (1989).

30. Wensink, H. H. et al. Meso-scale turbulence in living fluids. Proc Natl Acad Sci U S A 109, 14308–13 (2012).

31. Semmler, A. B., Whitchurch, C. B. & Mattick, J. S. A re-examination of twitching motility in Pseudomonas aeruginosa. Microbiology (N Y) 145, 2863–73 (1999).

32. Wei, Y. et al. The population dynamics of bacteria in physically structured habitats and the adaptive virtue of random motility. Proc Natl Acad Sci U S A 108, 4047–52 (2011).

33. Jin, F., Conrad, J. C., Gibiansky, M. L. & Wong, G. C. L. Bacteria use type-IV pili to slingshot on surfaces. Proc Natl Acad Sci U S A 108, 12617–22 (2011).

34. Creppy, A., Praud, O., Druart, X., Kohnke, P. L. & Plouraboué, F. Turbulence of swarming sperm. Phys Rev E 92, 032722 (2015).

35. Hood, R. D. et al. A Type VI Secretion System of Pseudomonas aeruginosa Targets a Toxin to Bacteria. Cell Host Microbe 7, 25–37 (2010).

36. Russell, A. B., Peterson, S. B. & Mougous, J. D. Type VI secretion system effectors: Poisons with a purpose. Nature Reviews Microbiology vol. 12 137–148 Preprint at https://doi.org/10.1038/nrmicro3185 (2014).

37. Smith, W. P. J. et al. The evolution of tit-for-tat in bacteria via the type VI secretion system. Nat Commun 11, 1–11 (2020).

38. Kudryashev, M. et al. Structure of the Type VI Secretion System Contractile Sheath Article Structure of the Type VI Secretion System Contractile Sheath. Cell 160, 952–962 (2015).

39. Hartmann, R. et al. Quantitative image analysis of microbial communities with BiofilmQ. Nat Microbiol 6, 151–156 (2021).

40. Hartmann, R., van Teeseling, M. C. F., Thanbichler, M. & Drescher, K. BacStalk: A comprehensive and interactive image analysis software tool for bacterial cell biology. Mol Microbiol 114, 140–150 (2020).

41. Jeckel, H. & Drescher, K. Advances and opportunities in image analysis of bacterial cells and communities. FEMS Microbiol Rev 45, 1–14 (2021).

42. van Valen, D. A. et al. Deep Learning Automates the Quantitative Analysis of Individual Cells in Live-Cell Imaging Experiments. PLoS Comput Biol 12, e1005177 (2016).

43. O’Connor, O. M., Alnahhas, R. N., Lugagne, J. B. & Dunlop, M. J. DeLTA 2.0: A deep learning pipeline for quantifying single-cell spatial and temporal dynamics. PLoS Comput Biol 18, e1009797 (2022).

44. Cuny, A. P., Ponti, A., Kündig, T., Rudolf, F. & Stelling, J. Cell region fingerprints enable highly precise single-cell tracking and lineage reconstruction. Nat Methods 19, 1276–1285 (2022).

45. Panigrahi, S. et al. Misic, a general deep learning-based method for the high-throughput cell segmentation of complex bacterial communities. Elife 10, e65151 (2021).

46. Bertrand, J. J., West, J. T. & Engel, J. N. Genetic Analysis of the Regulation of Type IV Pilus Function by the Chp Chemosensory System of Pseudomonas aeruginosa. J Bacteriol 192, 994–1010 (2009).

47. Schindelin, J. et al. Fiji: an open-source platform for biological-image analysis. Nat Methods 9, 676–682 (2012).

48. Thévenaz, P., Ruttimann, U. E. & Unser, M. A pyramid approach to subpixel registration based on intensity. IEEE Transactions on Image Processing 7, 27–41 (1998).

