## Supplementary information for "Tracking bacteria at high density with FAST, the Feature-Assisted Segmenter/Tracker"

December 2, 2022

**4 Appendix A: Entropy of a uniformly sampled random vector** **18**

### 1 Glossary of symbols

- Object features:
  - $\boldsymbol{r}$: Centroid of object.
  - $\phi$: Orientation of object.
  - $L_M$: Semi-major axis of object.
  - $L_m$: Semi-minor axis of object.
  - $A$: Area of object.
  - $m_c$: Mean intensity of object in channel $c$.
  - $s_c$: Standard deviation of object in channel $c$.
- Feature space notation:
  - $\boldsymbol{x}_t^i$: Feature vector constructed from any combination of the features listed above, associated with object $i$ in frame $t$.
  - $\{\boldsymbol{x}_t\}$: Set of feature vectors of objects detected in frame $t$.
  - $\bar{\boldsymbol{x}}_t^i$: Scaled version of $\boldsymbol{x}_t^i$.
  - $\hat{\boldsymbol{x}}_t^i$: Normalised version of $\boldsymbol{x}_t^i$.
  - $\{\Delta \boldsymbol{x}_t\}$: Set of displacements of objects in feature space between frames $t$ and $t+1$.
- User-selectable parameters:
  - $N$: Number of components of $\boldsymbol{x}_t^i$ (dimensionality of the feature space).
  - $F$: Fraction of links to be included in training dataset.
  - $P$: Ambiguous link probability.
  - $G$: Maximum bridgeable gap size.
- $Q_t^{i,j}$: Euclidean distance between feature vectors $\boldsymbol{x}_t^i$ and $\boldsymbol{x}_{t+1}^j$.
- $q_t^i$: Minimal Euclidean distance between $\boldsymbol{x}_t^i$ and the set of feature vectors $\{\boldsymbol{x}_{t+1}\}$.
- $k$: Denotes a linked pair of objects $\{i, j\}$ in frames $t$ and $t+1$ respectively.
- $n_{k,t}$: Number of training links made between objects in frame $t$ and objects in frame $t+1$.
- $n_{o,t}$: Number of objects present in frame $t$.
- $\boldsymbol{\mu}_t(\Delta \boldsymbol{x})$: Mean vector of $\{\Delta \boldsymbol{x}_t\}$.
- $\Sigma_t(\boldsymbol{x})$: Covariance matrix of $\{\boldsymbol{x}_t\}$.
- $\Sigma_t(\Delta \boldsymbol{x})$: Covariance matrix of $\{\Delta \boldsymbol{x}_t\}$.
- $I(X;Y)$: Mutual information between random variables $X$ and $Y$.
- $f(x,y)$: Joint probability density function (PDF) between random variables $X$ and $Y$.
- $f(x)$: Marginal PDF of random variable $X$.
- $H(X)$: Entropy of random variable $X$.
- $r_t$: Trackability score for frame $t$.
- $\beta_t$: Adaptive linking threshold for frame $t$.
- $g$: Current size of gap (number of frames) between compared timepoints.

#### 2 Software design goals

Following an extensive review of existing cell tracking software packages [1–6] and testing them on images of densely packed bacterial monolayers, we established several design criteria for our software:

1. Modular construction: Many existing tracking software packages use single GUI that contains all of their functionality. While this approach offers some advantages, the complexity of the resulting interface can be daunting to the novice user. Instead of this integrated approach, we decided to split FAST into several distinct modules, each with its own layout and workflow. The unique layout of each module within FAST allows users to visualise the results of each stage and directly optimise each user-defined parameter at intermediate points within the analysis, rather than having to rely solely on the final output.

2. Rapid feedback: Many software packages require the entire dataset to be analysed before the quality of analysis can be assessed. This can render the manual optimisation of analytical parameters impractical, as large datasets can take hours or even days to process. One of the critical goals of FAST was therefore to allow users to quickly analyse small subsets of their data and to provide immediate visual feedback on these analyses. Users can use these capabilities to rapidly optimise parameter sets. We also provide tools to rapidly identify portions of a timeseries where the tracking accuracy is diminished, allowing users to remove them from further analysis to reduce wasted computational effort.

3. Minimization of the number of user-defined parameters: As more and more features are incorporated into a tracking algorithm, the number of analytical parameters can become too large to optimise manually using an iterative approach. To avoid this, our machine learning-based tracking optimally integrates information across an arbitrary number of features. By calculating the statistical properties of each feature and automatically normalising them relative to one another, the user only needs to optimise a single tracking parameter.

4. Extensibility: Although many of the existing software packages we tested are powerful, in many cases they were designed to perform a specific task (e.g. lineage reconstruction [3] or measurement of extracellular structures [7,8]). FAST provides an extensible framework to test diverse hypotheses by facilitating the measurement of novel cell features and allowing custom analyses to be integrated into its pipeline. New features can be specified by users via a short script, while the results of novel custom analyses can easily be imported for visualisation with FAST. For example, users can use custom features (like cell shape or the localization pattern of a particular fluorescent protein) to assign trajectories to distinct populations (representing, for example, different bacterial species).

These overall design goals were applied to construct FAST, the architecture of which is shown in Fig. 1. In the following sections, we provide more detailed specifications of each of FAST's component modules, demonstrating how these aims were achieved.

#### 3 Module specifications

##### 3.1 Data import

Raw brightfield, phase contrast and fluorescence microscopy images can be imported into FAST from their native formats using Bioformats [9]. This supports a wide variety of common imaging formats, including the .czi, .nd2 and .lif files generated by commercial microscopy packages. The user can then choose to upsample the images by a factor of two, which can be useful for

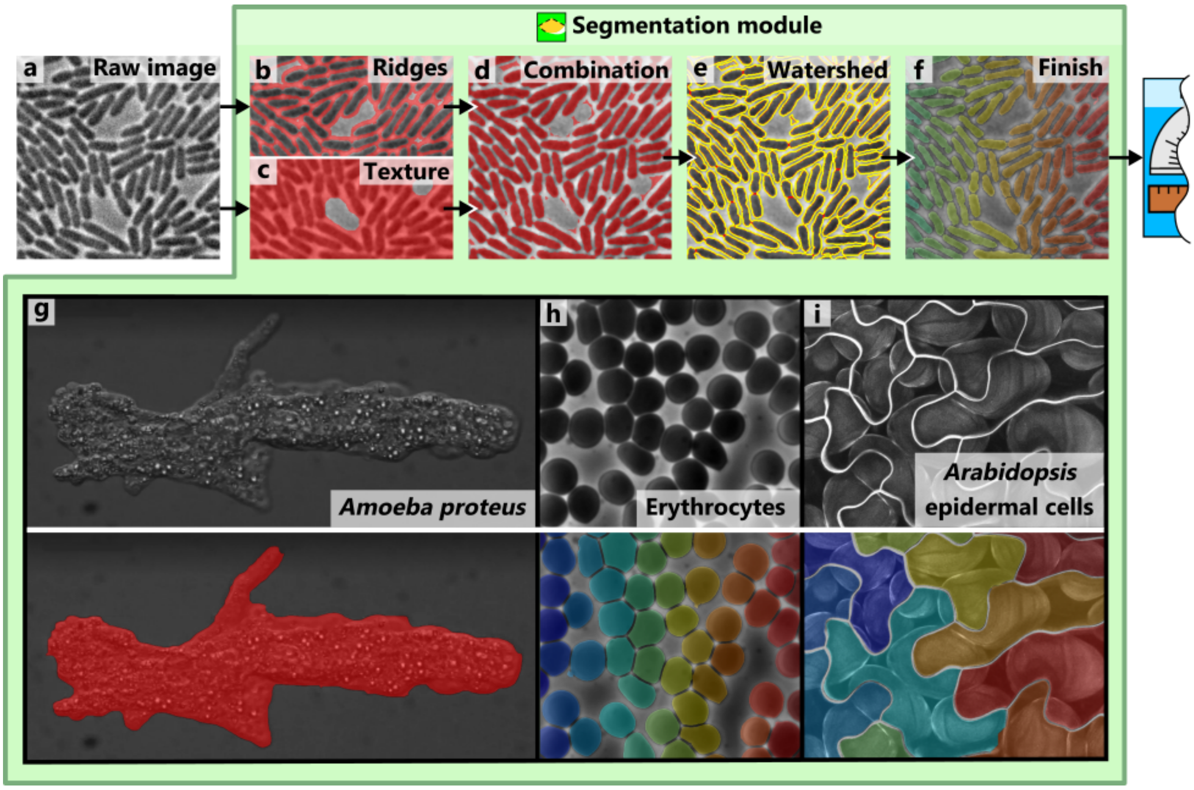

**Figure S1:** The Segmentation module. The segmentation pipeline is applied to a single channel within the dataset, typically the brightfield or phase-contrast channel (a). In the first stage, ridge detection (b) and texture analysis (c) are applied to the image, and are then combined to produce an approximate segmentation (d). This initial segmentation is then refined by using a morphological watershed algorithm to separate objects joined by small numbers of pixels (e). Finally, a filter based on the size of detected objects is applied, resulting in the final segmentation (f). The overlays shown in panels b-f are representative of the different types of visual feedback available within FAST's GUI, allowing fine-tuning of the parameters for each stage of the algorithm independently. Through the selection of appropriate parameters, microscopic samples with a wide variety of different properties can be accurately segmented (g-i) [11–13].

improving the quality of segmentations if individual objects are represented by a relatively small number of pixels.

To streamline later processing stages and reduce RAM requirements, FAST stores individual frames in each channel as separate .tif files in a directory structure. If data storage is a limiting factor during processing, these frames can be deleted after all analysis has been completed.

##### 3.2 The Segmentation module

The first step of the image analysis section of FAST generates a segmentation of one of the imaging channels, typically the brightfield or phase contrast (figure S1a). Following an initial pass with a median filter to reduce pixel noise, several methods are combined to accurately segment images with a wide variety of different properties. First, ridge detection [10] is applied to the image (figure S1b), which generates an outline of each object. As its name suggests, this algorithm is particularly effective at distinguishing closely-packed objects, separated by only a narrow, low-contrast ridge of pixels. Users can specify three parameters at this stage: the spatial scale of the ridges to be detected, the threshold used to convert the ridge score into a binary segmentation, and the minimum area of these segmented ridges.

In parallel, FAST applies a texture metric to the image. A box of user-specified size is drawn around each pixel location and the mean and standard deviation of the pixels within this box is measured. Dividing the standard deviation by the mean generates the coefficient of variation for the region around the target pixel. This is then multiplied by the square root of the mean intensity to account for differences in shot noise. The resulting metric is highly robust to changes in overall intensity caused by, for example, photobleaching of the sample. This is then converted into a second binary segmentation using another user-specified threshold, providing a coarse-grained separation of object-free regions from regions containing objects (figure S1c).

The results of these two segmentations are then combined using a pixelwise OR filter (figure S1d), followed by a morphological watershed segmentation step [14] (figure S1e). This allows the separation of dividing cells that are still partially attached to one another, as well as any objects not fully separated by the ridge detection algorithm. Finally, a pair of area filters are applied, allowing the user to reject objects that are too large or too small (figure S1f).

This combination of analytical stages can successfully segment a wide range of biological images, including low-contrast, densely-packed and morphologically complex objects (figure S1 g-i). Our pixel-based approach does not require direct user input for each object, unlike methods such as those that use deformable contours [15], nor does it constrain objects to conform to a specific morphological model, unlike many model-driven approaches (*e.g.* [16]). While these properties do mean that the occasional mis-segmentation is allowed to pass into later processing stages, this disadvantage is outweighed by the segmentation algorithm's speed and flexibility. As will be described in later sections, these mis-segmentations can be detected and removed during the tracking phase of the pipeline (section 3.4.5).

##### 3.3 The Feature Extraction module

The second stage of the pipeline, the Feature Extraction module, takes previously extracted segmentations and measures the features of associated objects (figure S2). The term 'feature' is generic and can in principle be applied to any quantitative property of an object, however here it is constrained to object centroid ($\boldsymbol{r}$), length $L_M$, width $L_m$, area $A$, orientation $\phi$, and the means $m_c$ and standard deviations $s_c$ of the pixel intensities associated with the object in each imaging channel $c$.

Two options are provided for measuring the morphological features $L_M$, $L_m$ and $\phi$. The 'ellipse fitting' choice applies an ellipse fitting routine to the object's boundary pixels and extracts these features as the parameters of the result. Alternatively, the 'perimeter distances' algorithm defines $L_M$ as the maximal distance between points on the boundary, and $\phi$ as the orientation of the line joining these two maximally separated points. Unwrapping the sequence of pixel locations defining the boundary to form an ordered list, $L_m$ is further defined as the distance between the pair of points on the boundary that are a) minimally separated in space and b) at least one quarter of the total boundary length away from each other in this ordered list (this constraint ensures that the selected points are on approximately opposite sides of the object). Ellipse fitting is most useful when objects have an approximately elliptical shape, as making use of all the points along the object's boundary renders it robust to noise. However, this approach can substantially overestimate the length of some long, thin objects - in these cases, the perimeter distances algorithm should be used instead.

To measure $m_c$ and $s_c$, the object's segmentation is first applied as a mask, defining a set of pixels in each imaging channel for which these statistics are calculated. Additionally, object area is measured as the total number of pixels in the object's mask.

Following measurement, these features are stored as a feature vector $\boldsymbol{x}_t^i$ for object $i$ in frame $t$. The number of components of this vector, $N$, represents the dimensionality of the feature space. In subsequent sections, curly braces will be used to denote sets of these feature vectors. For example, $\{\boldsymbol{x}_t\}$ represents the set of all feature vectors for all objects in frame $t$.

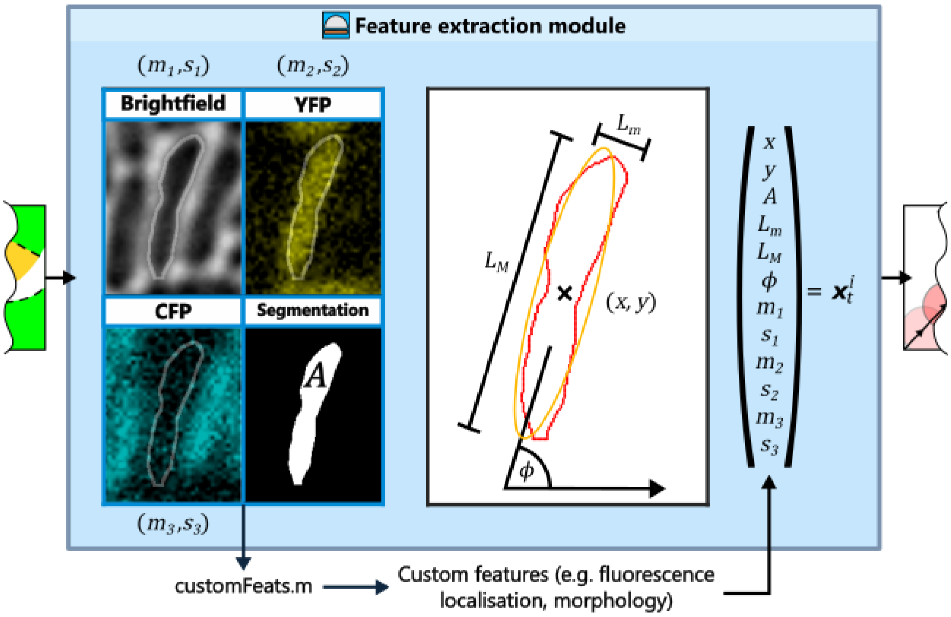

**Figure S2:** The Feature Extraction module. Users can choose to extract a number of commonly used features from objects, including the mean $m$ and standard deviation $s$ of each of the input channels (here brightfield, YFP and CFP), the area $A$, length $L_M$ and width $L_m$ of the object, as well as its position $(x, y)$ and orientation $\phi$ with respect to the fixed image frame. Novel, user-defined features can also be extracted from a combination of the raw images and segmentation through the customFeats.m function. The set of measured features forms the feature vector $\boldsymbol{x}_t^i$ associated with object $i$ in frame $t$.

##### 3.3.1 Custom features

The Feature Extraction module's default set of features has been chosen as the most generally useful for addressing a wide range of questions. However, some users may wish to measure other quantities, such as the distribution of fluorescence along the length of cells or morphological measures more sophisticated than $L_M$ and $L_m$. To facilitate this, FAST supports the definition of custom features using the 'customFeats.m' function. This is passed images of each object in each available imaging channel from the Feature Extraction module, along with the associated segmentation mask and pixel size. Users can manipulate these inputs to extract feature information, which is then returned to the Feature Extraction module. Custom features extracted in this way are incorporated directly into the feature vector $\boldsymbol{x}_t^i$, and are thereby integrated into downstream analyses.

An example application of this functionality is given in Fig. 6b, with more detailed documentation (including a function template) available on the project wiki (`https://bit.ly/3gnBcCY`).

#### 3.4 The Tracking module

Having extracted the feature vector $\boldsymbol{x}_t^i$ associated with each object, the problem of tracking becomes one of assigning links between objects in subsequent timepoints through comparisons of $\{\boldsymbol{x}_t\}$ with $\{\boldsymbol{x}_{t+1}\}$. At its heart, this is a distance-measuring problem, where we wish to establish links between objects in frames $t$ and $t + 1$ that are close to each other in feature space. Indeed, the most basic tracking algorithm is simply to measure the positional distance between all pairs of objects in the two frames and link together neighbouring pairs.

Unfortunately, there are several ways in which such a simple nearest-neighbour approach can generate spurious results. Firstly, objects can spontaneously move into and out of the field of view, requiring new tracks to be initiated and existing tracks to be terminated, respectively.

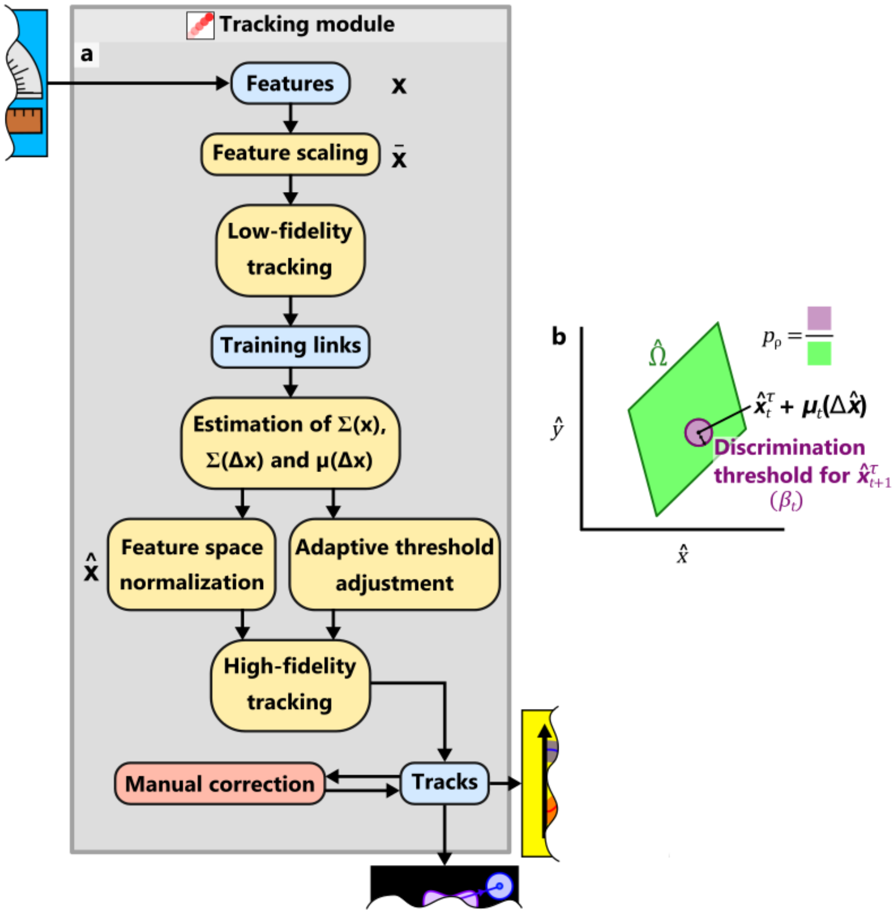

**Figure S3:** The Tracking module. a) Flow diagram indicating the stages of FAST's tracking algorithm. The stages at which the feature vectors $\boldsymbol{x}$, scaled feature vectors $\bar{\boldsymbol{x}}$ and normalised feature vectors $\hat{\boldsymbol{x}}$ are used are indicated. b) Following feature normalization, $\hat{\Omega}$ – the support of the distribution $f(\hat{\boldsymbol{x}})$ – can in general be represented as a parallelepiped. $p_\rho$ is the probability that a random object $\rho$ drawn uniformly from this support appears within the discrimination threshold of a given target $\tau$ (purple shaded area), as opposed to elsewhere within the support (green shaded area).

Secondly, objects can be mis-segmented - split into more than one segmented region or not detected at all - in frame $t+1$, meaning links for the corresponding objects in frame $t$ should be suppressed for this timepoint. Thirdly, noise or unrelated objects ('clutter') may be mistaken for the target.

The core aim of FAST's Tracking module is to reject these false links optimally, while retaining as many true links as possible. Our approach is based on the assumption that, on average, each object's appearance/position/orientation etc (the feature data stored in $\boldsymbol{x}_t^i$) either remains stable or changes predictably from one frame to the next. Discontinuities in feature data are associated with false links - a large apparent change in object area combined with a moderate change in object position, for example, would be indicative of mis-segmentation.

Of course, not all features are equally useful in this regard. Our ability to predict the value of a particular feature in frame $t+1$ based on its value in frame $t$ depends on the amount of noise occurring between frames, with some features being more noisy than others. A key step on the way to this optimised link rejection, therefore, is to automatically determine how different features should be weighted relative to one another when making comparisons between objects in sequential frames. We also wish to adjust how stringently we filter proposed links on the basis of the total amount of noise present in the dataset, as noisier datasets put us at risk of making a greater number of false links.

The overarching framework that we use to achieve these goals is information theory, a field of statistics concerned with how our ability to predict a particular outcome changes on the basis of prior observations. A typical example is to ask how well can we predict the contents of a complete message based on only a fragment of the whole. If circumstances allow us to predict with certainty the contents of the entire transmission from a subsection, we can deduce that the unseen remainder of the message contains no additional information. At first glance, it may not seem that this type of question has much applicability to object tracking, but it turns out that tackling a very similar question - how much does our certainty about the position of an object in frame $t+1$ increase as a result of the observation of its position in frame $t$? - provides us with an elegant solution to the problems stated above.

The FAST tracking algorithm consists of three main stages (figure S3a):

- Measurement of the ease of tracking (equivalent to the amount of information available) at each timepoint.
- Rescaling of features according to the amount of information they contribute to solving the linking problem.
- Calculation of a distance-based discrimination threshold that suppresses links when there is insufficient information for them to be unambiguously assigned.

These stages will be described in more detail in sections 3.4.2 - 3.4.4. Fundamental to all three, however, is the generation of a training dataset from which the time-dependent statistics necessary for subsequent stages can be extracted. This forms the first stage of the tracking algorithm.

##### 3.4.1 Generating the training dataset

The training dataset is generated from the input feature data by finding the easiest to assign links using a simple Euclidean distance-based tracking algorithm. To provide some degree of balance between different features during this first tracking run, feature scaling is applied [17]. Defining the maximum value of each feature in each frame to be $\max(\{\boldsymbol{x}_t\})$ and the minimum value to be $\min(\{\boldsymbol{x}_t\})$, the range of all features is set to $[0, 1]$ using the equation:

$$\bar{\boldsymbol{x}}_t^i = \frac{\boldsymbol{x}_t^i - \min(\{\boldsymbol{x}_t\})}{\max(\{\boldsymbol{x}_t\}) - \min(\{\boldsymbol{x}_t\})}. \tag{1}$$

A list of putative links between objects in frame $t$ and $t+1$ is then created, pairing objects according to the minimal Euclidean distance between $\bar{\boldsymbol{x}}_t^i$ and $\bar{\boldsymbol{x}}_{t+1}^j$. To be explicit, the matrix $Q_t^{i,j}$ of Euclidean distances between objects in sequential frames is calculated as

$$Q_t^{i,j} = \|\bar{\boldsymbol{x}}_{t+1}^j - \bar{\boldsymbol{x}}_t^i\|, \tag{2}$$

where $\|\cdot\|$ indicates the Euclidean norm of the contents. Next, the unique score $q_i$ is found that satisfies:

$$q_i = \min_j(Q_t^{i,j}). \tag{3}$$

The links with scores $q_i$ are then sorted by score, and the proportion, $F$, of lowest scoring links classified as correct. $F$ is the first user-adjustable parameter of the algorithm.

One problem with this approach is that equation 1 tends to weight low-quality and high-quality features equally. This can severely limit the accuracy of links based on the simple similarity metric $q_i$. To avoid this problem, the user is provided with the option of restricting the scaled vector $\bar{\boldsymbol{x}}_t^i$ to the coordinates of the object's centroid, $\bar{\boldsymbol{r}}_t^i$. This is typically robust enough for this initial, relatively low-fidelity tracking stage; nevertheless, some datasets with large amounts of motion between frames and high object densities may in some cases require use of the full complement of features.

From this set of initial training links, we can extract the frame-frame feature displacements for each link $k$, $\Delta \boldsymbol{x}_t^k = \boldsymbol{x}_{t+1}^k - \boldsymbol{x}_t^k$. The set of these displacement measurements at each timepoint, $\{\Delta \boldsymbol{x}_t\}$, contains the information necessary to measure the predictable and unpredictable components of each object's motion through feature space. Specifically, we calculate the mean vector representing the average change in each feature between frames, $\boldsymbol{\mu}_t(\Delta \boldsymbol{x})$, as:

$$\boldsymbol{\mu}_t(\Delta \boldsymbol{x}) = \frac{1}{n_{k,t}} \sum_{k=1}^{n_{k,t}} \Delta \boldsymbol{x}_t^k, \tag{4}$$

where $n_{k,t}$ is the total number of links in the training dataset. We also measure the extent of correlations between objects' movements in feature space by calculating the sample covariance matrix $\Sigma_t(\Delta \boldsymbol{x})$ from our training dataset:

$$\Sigma_t(\Delta \boldsymbol{x}) = \frac{1}{n_{k,t} - 1} \sum_{k=1}^{n_{k,t}} (\Delta \boldsymbol{x}_t^k - \boldsymbol{\mu}_t(\Delta \boldsymbol{x}))(\Delta \boldsymbol{x}_t^k - \boldsymbol{\mu}_t(\Delta \boldsymbol{x}))^\intercal. \tag{5}$$

We can also calculate the equivalent statistics $\boldsymbol{\mu}_t(\boldsymbol{x})$ and $\Sigma_t(\boldsymbol{x})$ for the instantaneous object locations in feature space by substituting the raw feature vectors $\{\boldsymbol{x}_t\}$ in place of the feature displacements $\{\Delta \boldsymbol{x}_t\}$ in equations 4 and 5.

##### 3.4.2 Quantification of dataset information

One goal of the FAST tracking algorithm is to provide a robust metric of projected tracking accuracy based on the limits of the feature information available within the dataset. Qualitatively, we can see that the usefulness of each feature for assigning object identity between frames will depend on two factors (Fig. 2a-c): firstly, the stability of the feature between frames (with more noisy features being less reliable), and secondly, the dynamic range of the feature (since a larger range of values will allow a larger number of objects to be distinguishable for a given amount of noise). Our metric should therefore somehow capture the contribution of these two factors to the link assignment process, and integrate these measures across all features.

Information theory provides us with a natural definition for this metric: the mutual information between successive frames. We assume that our objects can be found at an initial location in feature space $\boldsymbol{x}_t$, and subsequently move to a different location, $\boldsymbol{x}_{t+1}$. Treating these positions as random vectors $\boldsymbol{X}_t$ and $\boldsymbol{X}_{t+1}$, we can calculate the mutual information $I(\boldsymbol{X}_t; \boldsymbol{X}_{t+1})$ between them as [18]

$$I(\boldsymbol{X}_t; \boldsymbol{X}_{t+1}) = \int_{-\infty}^{\infty} \int_{-\infty}^{\infty} f(\boldsymbol{x}_t, \boldsymbol{x}_{t+1}) \log_2 \frac{f(\boldsymbol{x}_t, \boldsymbol{x}_{t+1})}{f(\boldsymbol{x}_t) f(\boldsymbol{x}_{t+1})} \, d\boldsymbol{x}_t \, d\boldsymbol{x}_{t+1}, \tag{6}$$

where $f(\boldsymbol{x}_t)$ indicates the probability density function (PDF) of $\boldsymbol{X}_t$, and $f(\boldsymbol{x}_t, \boldsymbol{x}_{t+1})$ indicates the joint PDF between $\boldsymbol{X}_t$ and $\boldsymbol{X}_{t+1}$. $I(\boldsymbol{X}_t; \boldsymbol{X}_{t+1})$ provides a precise measure of the increase of our certainty about the value of $\boldsymbol{x}_{t+1}$, given our observation of $\boldsymbol{x}_t$ at the prior timepoint and knowledge of the relationship between $\boldsymbol{X}_t$ and $\boldsymbol{X}_{t+1}$.

In principle, we could evaluate equation 6 at each timepoint using, for example, histogram-based methods to estimate $f(\boldsymbol{x}_t)$, $f(\boldsymbol{x}_{t+1})$ and $f(\boldsymbol{x}_t, \boldsymbol{x}_{t+1})$. However, to measure variations of the information content of the dataset over time we would need to evaluate these terms at each timepoint. In practice, the sparseness of the resulting data means that these estimated distributions are usually insufficiently robust for accurate measurements of $I(\boldsymbol{X}_t; \boldsymbol{X}_{t+1})$. Instead, we make use of the alternative formulation for the mutual information [18]:

$$I(\boldsymbol{X}_t; \boldsymbol{X}_{t+1}) = H(\boldsymbol{X}_{t+1}) - H(\boldsymbol{X}_{t+1}|\boldsymbol{X}_t), \tag{7}$$

where $H(X)$ indicates the entropy of $X$, and is given by:

$$H(X) = -\int_{-\infty}^{\infty} f(x)\log_2(f(x)) \cdot dx, \tag{8}$$

and $H(\boldsymbol{X}_{t+1}|\boldsymbol{X}_t)$ indicates the conditional entropy of $\boldsymbol{X}_{t+1}$ at time $t+1$ given knowledge of the value of $\boldsymbol{X}_t$ at time $t$.

We can easily evaluate equation 7 by assuming that $f(\boldsymbol{x}_t)$ and $f(\boldsymbol{x}_{t+1}|\boldsymbol{x}_t)$ can be approximated by analytically tractable distributions, the parameters for which we can extract from the training dataset (for practical and notational purposes, we assume that $f(\boldsymbol{x}_t) = f(\boldsymbol{x}_{t+1})$). In this case, we assume that $\boldsymbol{x}_t^i$ is drawn with equal probability (*i.e.* uniformly) from a fixed region of feature space. To be precise,

$$f(\boldsymbol{x}_t) = \begin{cases} (\int_\Omega 1 \cdot d\boldsymbol{x}_t)^{-1} & \text{for} \quad \boldsymbol{x}_t \in \Omega \\ 0 & \text{for} \quad \boldsymbol{x}_t \notin \Omega, \end{cases} \tag{9}$$

where $\Omega$ represents the support of the distribution, the region of feature space where an object might be found. For simplicity, we assume here that this region is shaped as a parallelepiped, which simultaneously allows accurate representation of the full range of possible centroid positions (assuming objects are uniformly distributed throughout the image) while still capturing possible correlations between other features. This distribution is analagous to a one-dimensional Uniform distribution in that it is constant over its support, though its ability to also capture correlations between features provides an additional level of flexibility.

The boundaries of this parallelepiped can be represented by the matrix $(12\Sigma_t(\boldsymbol{x}))^{\frac{1}{2}}$, where $\Sigma_t(\boldsymbol{x})$ is the covariance matrix of $\{\boldsymbol{x}_t\}$ as calculated in section 3.4.1. Because the entropy of a uniformly sampled random vector is simply equal to the log of the integral of its support (appendix A), we can calculate the entropy $H(\boldsymbol{X}_t)$ by measuring the (hyper)volume of this parallelepiped:

$$\begin{aligned} H(\boldsymbol{X}_t) &= \log_2 |(12\Sigma_t(\boldsymbol{x}))^{\frac{1}{2}}| \\ &= \log_2(12^{\frac{N}{2}}|\Sigma_t(\boldsymbol{x})^{\frac{1}{2}}|) \\ &= \frac{1}{2}\log_2(|\Sigma_t(\boldsymbol{x})|) + \frac{N}{2}\log_2(12), \end{aligned} \tag{10}$$

where $|\cdot|$ indicates the determinant of the contents.

Similarly, to represent the stochastic motion of the object through feature space, we can assume that $f(\boldsymbol{x}_{t+1}|\boldsymbol{x}_t)$ is drawn from a multivariate Gaussian distribution, $\mathcal{N}(\boldsymbol{\mu}_t(\Delta\boldsymbol{x}), \Sigma_t(\Delta\boldsymbol{x}))$, where $\boldsymbol{\mu}_t(\Delta\boldsymbol{x})$ and $\Sigma_t(\Delta\boldsymbol{x})$ are statistics calculated in section 3.4.1. The mutual information between the random vectors $\boldsymbol{X}_t$ and $\boldsymbol{X}_{t+1}$ can now be calculated by substituting the standard expression for the entropy of the multivariate normal [19] and equation 10 into equation 7:

$$I(\boldsymbol{X}_t; \boldsymbol{X}_{t+1}) = \frac{1}{2}\log_2\left(\frac{|\Sigma_t(\boldsymbol{x})|}{|\Sigma_t(\Delta\boldsymbol{x})|}\right) + \frac{N}{2}\left(\log_2\left(\frac{6}{\pi e}\right)\right). \tag{11}$$

Our final metric of the ease of tracking objects between frames $t$ and $t+1$, referred to as the trackability score $r_t$, also takes into account the total number of objects present at this timepoint:

$$r_t = I(\boldsymbol{X}_t; \boldsymbol{X}_{t+1}) - \log_2(n_{o,t}), \tag{12}$$

where $n_{o,t}$ indicates the number of objects present at time $t$.

$r_t$ indicates the average amount of information available for distinguishing each object in frame $t+1$ from one another. Relatively small values of $r_t$ correspond to situations where the amount of unpredictable noise during an object's movement through feature space is sufficiently large for there to be several plausible candidate links; in these ambiguous cases, accurate tracking is generally impossible unless additional information can be added (through, for example, the addition of extra features) or the number of objects reduced. As shown in Fig. 4a, $r_t$ can also be used to determine when problems have occurred during imaging. In principle, this may allow it to be used as a means of automatically flagging and/or expunging problematic sections of datasets.

##### 3.4.3 Feature normalisation

Although we can now measure how much information is available in our dataset at each timepoint, it has still not been established how to optimally make use of that information. When generating the training dataset, we made use of feature scaling to combine measurements from multiple different features (equation 1). This is appropriate for the comparatively low-fidelity tracking necessary for generating the training dataset, as the summary statistics $\boldsymbol{\mu}_t(\Delta\boldsymbol{x})$ and $\Sigma_t(\Delta\boldsymbol{x})$ are robust to the omission of the most difficult to assign links. For performing high-fidelity tracking however, it is clearly inappropriate: firstly, it weights both high-information and low-information features equally, and secondly it ignores possible correlations between features. In addition, predictable changes in feature values over time (for example, the gradual increase in the length of cells due to growth) are neglected.

To resolve these problems, FAST employs a technique we call feature normalisation. In geometrical terms, the feature space is shifted, scaled and rotated such that the amount of noise that each transformed feature contributes is equal and zero-centred (Fig. 2d). This approach is similar to that described in [20], although importantly our use of the initial training dataset to collect the necessary summary statistics allows this process to be automated, rather than requiring laborious manual collection as previously described.

Mathematically, we can describe the necessary transformation to convert the feature space $\boldsymbol{x}_t$ to the normalised displacement space $\Delta\hat{\boldsymbol{x}}_t$ as:

$$\Delta\hat{\boldsymbol{x}}_t^{i,j} = \Sigma_t(\Delta\boldsymbol{x})^{-\frac{1}{2}}(\boldsymbol{x}_{t+1}^j - \boldsymbol{x}_t^i - \boldsymbol{\mu}_t(\Delta\boldsymbol{x})). \tag{13}$$

This equation is the normalised equivalent to equation 2, prior to the application of the Euclidean norm. Here, the term $\Sigma_t(\Delta\boldsymbol{x})^{-\frac{1}{2}}$ indicates the inverse of the principal square root of the covariance matrix $\Sigma_t(\Delta\boldsymbol{x})$. This term ensures that the covariance matrix of the transformed set of feature vectors is equal to the $N$-dimensional identity matrix. For this reason it can also be described as a whitening transformation, as it removes correlations between feature displacements. Combined with the subtraction of the mean vector $\boldsymbol{\mu}_t(\Delta\boldsymbol{x})$, this transformation ensures that the normalised set of displacement vectors $\{\Delta\hat{\boldsymbol{x}}_t\}$ for the true links is isotropic and zero-centred, as required. FAST allows direct visualisation of $\{\Delta\hat{\boldsymbol{x}}_t\}$, permitting the user to compare pairs of normalised feature displacements and so assess whether the model has been correctly trained.

This calculation is the multivariate extension of the calculation of the standard score for a univariate normal distribution, and results in a metric known as the Mahalanobis distance. By analogy, just as we would trust a univariate measurement that was one standard deviation away from its predicted location over one that was three standard deviations away, so we can use this transformed distance to distinguish reliable links (small $\Delta\hat{\boldsymbol{x}}_t^{i,j}$) from false links arising from clutter and mis-segmentations (large $\Delta\hat{\boldsymbol{x}}_t^{i,j}$). Our robust tracking stage proceeds by calculating the normalised distance matrix $\hat{Q}_t^{i,j}$ using this metric, substituting the term $\bar{\boldsymbol{x}}_{t+1}^j - \bar{\boldsymbol{x}}_t^i$ with the normalised displacements $\Delta\hat{\boldsymbol{x}}_t^{i,j}$ in equation 2. Also similar to the method used to generate the training dataset, we keep putative links that minimise this distance, generating a score associated with each proposed link of $\hat{q}_i$ (equation 3). However, instead of choosing the lowest scoring fraction $F$ of putative links as during the generation of the training dataset, we only keep links which have a score $\hat{q}_i$ below the distance threshold $\beta_t$.

##### 3.4.4 The adaptive tracking threshold

As discussed in the opening to this section, the core objective of FAST's tracking algorithm is to optimally distinguish true links from false links. Given this objective, it makes sense to vary the stringency of the distance threshold $\beta_t$ based on the probability of false links occurring: when we are at high risk of making a false link, we should use a small value of $\beta_t$, maintaining only those links for which we have the greatest amount of evidence. In contrast, when the risk of making a false link is low, we can allow $\beta_t$ to increase, expanding the detection region and capturing a greater number of the true links in the dataset. FAST therefore allows the user to select a parameter that approximates the false link probability for each link, and automatically varies the value of $\beta_t$ to match.

Unfortunately, directly calculating the rate at which false links occur is challenging, as it requires us to know the probability of the different types of false link (links to mis-segmented objects, links to clutter and links to objects that have just moved into/out of frame). Instead, we consider a user-selected ambiguous link probability $P$, which is the probability that an object $\rho$ unrelated to the target $\tau$ is located within the radius $\beta_t$ (figure S3b). If $\tau$ disappears in the next frame - either because it moves out of frame or is mis-segmentated - $\rho$ will appear sufficiently close in appearance and position to be mistaken for $\tau$. On the other hand, if $\rho$ is simply random clutter, its appearance within this radius means it still has a chance of being linked preferentially over $\tau$, provided it is closer than $\tau$ to $\tau$'s predicted location. Thus, this condition - that a random object is found within the region defined by $\beta_t$ - must be met for any of the three types of false link to occur. This ambiguous link probability is therefore an appropriate quantity on which to base automated adjustments to the value of $\beta_t$.

To calculate the relationship between $P$ and $\beta_t$, we begin by assuming that the feature vector $\boldsymbol{x}_{t+1}^{\rho}$ of a single unrelated object $\rho$ is sampled uniformly from the distribution $f(\boldsymbol{x}_t)$, just as we assumed for the feature vector $\boldsymbol{x}_t^{\tau}$ of the target $\tau$ at the previous timepoint. This vector is then normalised in the same way as the target (equation 13), resulting in it being a distance of $\Delta\hat{\boldsymbol{x}}_t^{\tau,\rho}$ standard deviations away from the predicted location of $\tau$.

The distribution $f(\hat{\boldsymbol{x}}_t)$ is similar to the previously discussed distribution $f(\boldsymbol{x}_t)$, in that the sets of normalised feature vectors $\{\hat{\boldsymbol{x}}_t\}$ and $\{\hat{\boldsymbol{x}}_{t+1}\}$ are drawn uniformly from its support. We can derive its covariance matrix $\Sigma_t(\hat{\boldsymbol{x}})$ by applying the feature normalisation transformation to the covariance matrix of $f(\boldsymbol{x}_t)$, arriving at $\Sigma_t(\hat{\boldsymbol{x}}) = \Sigma_t(\Delta\boldsymbol{x})^{-1}\Sigma_t(\boldsymbol{x})$. We can therefore calculate the entropy of $\hat{\boldsymbol{X}}_t$ as:

$$H(\hat{\boldsymbol{X}}_t) = \frac{1}{2}\log_2\left(\frac{|\Sigma_t(\boldsymbol{x})|}{|\Sigma_t(\Delta\boldsymbol{x})|}\right) + \frac{N}{2}\log_2(12). \tag{14}$$

The distance threshold $\beta_t$ can be visualised as an $N$-sphere centred on $\boldsymbol{x}_t^i + \boldsymbol{\mu}_t(\Delta\boldsymbol{x})$ marking the boundaries of a sub-region of the support $\hat{\Omega}$ of $f(\hat{\boldsymbol{x}}_t)$ (figure S3b). To calculate the probability that our random object $\rho$ falls within this region (which is the ambiguous link probability $P$ when only $\tau$ and $\rho$ are present, so $n_{o,t} = 2$), we can find the self-information between $f(\hat{\boldsymbol{x}}_t)$ and the distribution $f(\hat{\boldsymbol{x}}_t^{\rho}|\Delta\hat{\boldsymbol{x}}_t^{\tau,\rho} \leq \beta_t)$. This latter PDF is the distribution of $\hat{\boldsymbol{X}}_t^{\rho}$ conditional on it being less than a distance $\beta_t$ away from the predicted location of $\tau$ (note that the Mahalanobis distance is again the metric of this space). This self-information can be interpreted as our degree of surprise at finding the object $\rho$ randomly inside the region set by $\beta_t$. Using the standard expression for the volume of an $N$-sphere, we can write this new PDF as:

$$f(\hat{\boldsymbol{x}}_t^{\rho}|\Delta\hat{\boldsymbol{x}}_t^{\tau,\rho} \leq \beta_t) = \begin{cases} \frac{\Gamma(1+N/2)}{(\beta_t\sqrt{\pi})^N} & \text{for} \quad \Delta\hat{\boldsymbol{x}}_t^{\tau,\rho} \leq \beta_t \\ 0 & \text{for} \quad \Delta\hat{\boldsymbol{x}}_t^{\tau,\rho} > \beta_t \end{cases} \tag{15}$$

where $\Gamma()$ indicates the gamma function. Using the expression for the entropy of uniformly sampled random variables (appendix A), we can now once again use equation 7 to measure the self-information $I_s$ as:

$$I_s = \frac{1}{2}\log_2\left(\frac{|\Sigma_t(\boldsymbol{x})|}{|\Sigma_t(\Delta\boldsymbol{x})|}\right) + \frac{N}{2}\log_2(12) - \log_2\left(\frac{(\beta_t\sqrt{\pi})^N}{\Gamma(1+N/2)}\right) \tag{16}$$

We can convert this into the probability, $p_\rho$, of observing a single unrelated object randomly inside this radius as

$$p_\rho = 2^{-I_s}. \tag{17}$$

Assuming that each of the $n_{o,t}-1$ objects unrelated to the target that are present at this time are sampled i.i.d. from $f(\hat{\boldsymbol{x}}_t)$, we can write $P$ as

$$P = 1 - (1-p_\rho)^{n_{o,t}-1}, \tag{18}$$

which, if $p_\rho \ll 1$, can be approximated as $P \approx p_\rho(n_{o,t}-1)$. Solving this set of equations for $\beta_t$, we can finally write down the desired threshold radius as a function of the user-selected value of $P$:

$$\beta_t = \sqrt[N]{P}\left(\frac{12}{\pi}\right)^{\frac{1}{2}}\left(\frac{\Gamma(1+N/2)}{(n_{o,t}-1)}\right)^{\frac{1}{N}}\left(\frac{|\Sigma_t(\boldsymbol{x})|}{|\Sigma_t(\Delta\boldsymbol{x})|}\right)^{\frac{1}{2N}}. \tag{19}$$

We again see the term $\frac{|\Sigma_t(\boldsymbol{x})|}{|\Sigma_t(\Delta\boldsymbol{x})|}$ appearing in this expression, suggesting a connection between $\beta_t$ and the information available to the tracking algorithm (equation 11). In brief, this connection arises because there is an inverse relationship between the amount of information available and the density of objects in the normalised feature space. When there is a large amount of information available, objects are widely spaced in the normalised feature space, and we can afford to search a large distance away from the predicted location of the target before we risk encountering an object that is not the target. In contrast, when the amount of available information is low, objects will be very densely packed together, and unrelated objects may appear close to the predicted location of the target. This expression therefore allows us to automatically and dynamically account for this relationship during tracking, helping to ensure that tracking accuracy remains consistent even if the statistical properties of the dataset vary widely over time.

##### 3.4.5 Gap bridging

This adaptive threshold $\beta_t$ will filter out many putative links, leaving objects that lack connections to the next frame. In some instances, such as when the target object moves out of the imaging region, trajectories should end at this point. In other cases though, link rejection corresponds to a temporary problem with object detection - for example, due to the mis-segmentation of an object in a single frame. In these cases, we wish to bridge the resulting gap between accurate segmentations.

To achieve this, FAST re-runs its tracking algorithm after its initial sweep through the dataset. However, instead making comparisons between $\hat{\boldsymbol{x}}_t^i$ and $\hat{\boldsymbol{x}}_{t+1}^j$ in equation 13, we now make comparisons $\hat{\boldsymbol{x}}_t^i$ and $\hat{\boldsymbol{x}}_{t+2}^j$. Objects linked from and to during the initial sweep through the dataset are removed from $\{\boldsymbol{x}_t\}$ and $\{\boldsymbol{x}_{t+2}\}$, preventing multiple assignments to single objects being made. Working under the assumption that objects move diffusively through feature space over time, $\beta_t$ is also rescaled by $\sqrt{2}$. This reduction in stringency allows us to account for the increasing displacement of objects from their initial positions during the period in which they go unobserved.

In general, we can repeat this process as many cycles as desired, incrementing the time gap $g$ by one frame each cycle such that $\hat{\boldsymbol{x}}_t^i$ is compared with $\hat{\boldsymbol{x}}_{t+g}^j$ and $\beta_t$ is rescaled by $\sqrt{g}$. The user selectable parameter $G$ represents the maximum value of $g$, and therefore corresponds to the maximum number of sequential frames that can be bridged as a single gap in a trajectory.

###### 3.4.6 Manual track correction

Regardless of the sophistication of the segmentation and tracking algorithms employed, automated methods will always generate some errors that are detectable by eye. It has been noted that in the drive for complete automation, many tools lack the option to manually adjust and correct tracks [21]. FAST enables this option through its validation GUI, which provides visual tools for identifying and breaking incorrectly assigned links, as well as assigning any links that were missed by the tracking algorithm.

Whether or not manual track correction is performed, FAST allows users to choose a minimum track length threshold to apply their final track data. This allows short tracks (typically corresponding to temporarily mis-segmented objects and noise) to be removed from downstream analyses.

##### 3.5 The Division Detection module

The tracking algorithm described in the previous sections can easily be adapted to permit detection of cellular divisions with several adjustments (figure S4a):

- In place of comparisons between the feature vectors of objects in adjacent frames, we instead need to compare $\{\boldsymbol{x}_m\}$ and $\{\boldsymbol{x}_d\}$, the feature vectors of mother and daughter cells respectively. Because tracks begin at cell birth and end at cell division, $\{\boldsymbol{x}_m\}$ and $\{\boldsymbol{x}_d\}$ can be compiled across all timepoints by finding the last (mother) and first (daughter) feature vectors associated with each track.

- We also make a prediction of the locations of the two daughter cells in feature space based on the mother's location before division (figure S4b). For example, in the case of the division of most bacteria, area and length typically halve, while the centroids of daughter cells can be inferred from the mother cell's centroid, length and orientation. These predicted positions $\boldsymbol{x}_p^i$ take the place of terms based on $\boldsymbol{x}_t^i$ in sections 3.4.1-3.4.4, while the actual positions $\boldsymbol{x}_d^j$ take the place of terms based on $\boldsymbol{x}_{t+1}^j$ (figure S4c).

- Because the track of the mother cell does not necessarily end at the same time as the daughter cell tracks begin (due to potentially unstable separation of daughter cells by the segmentation module), we append the end frame indices of the daughters' tracks to $\boldsymbol{x}_p^i$ and the mother's start frame index to $\boldsymbol{x}_d^j$. These frame indices are now simply treated as another feature, allowing us to automatically compensate for the variable stability of the separation of the two daughter cells by the segmentation algorithm. Gap bridging is performed directly by this process, preventing the need for user selection of $G$.

- As $\{\boldsymbol{x}_p\}$ and $\{\boldsymbol{x}_d\}$ are compiled across all timepoints, the time-dependence of dataset statistics (*e.g* the gradual bleaching of cell fluorescence over the course of an experiment) is no longer represented in the training data. As a result, only a single trackability score is calculated for the entire dataset, and the user is asked to select the distance threshold $\beta_t$ directly.

The remaining steps proceed exactly as described for cell tracking. After division detection has terminated, users can select a minimal lineage size threshold to filter the data that will be included in downstream analyses.

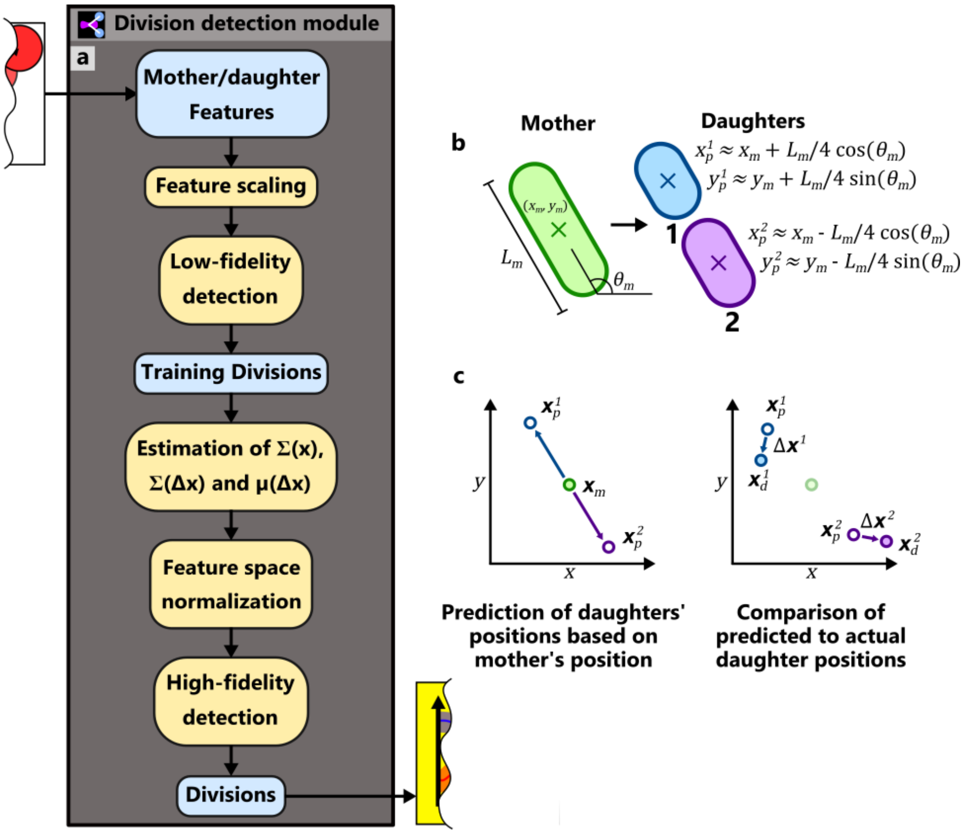

**Figure S4:** The Division Detection module. a) Flow diagram of the principal stages of the division detection process. Note that the division detection algorithm is similar to that described in figure S3a. b) The location of daughter cells in feature space can be usually be predicted from the location of the mother. In the example shown of a bacillus, the length of the two daughters can be predicted as half of that of the mother, while the $x$ and $y$ coordinates of the daughters can be predicted from the position, length and orientation of the mother. c) To adapt the tracking algorithm to perform division detection, we substitute the predicted positions of the daughters $\boldsymbol{x}_p^i$ in place of $\boldsymbol{x}_t^i$ in sections 3.4.1-3.4.4, and the actual positions of the daughters $\boldsymbol{x}_d^j$ in place of $\boldsymbol{x}_{t+1}^j$. The displacements $\Delta\boldsymbol{x}^k$ then correspond to the difference between the predicted positions of the daughters in feature space and their actual positions.

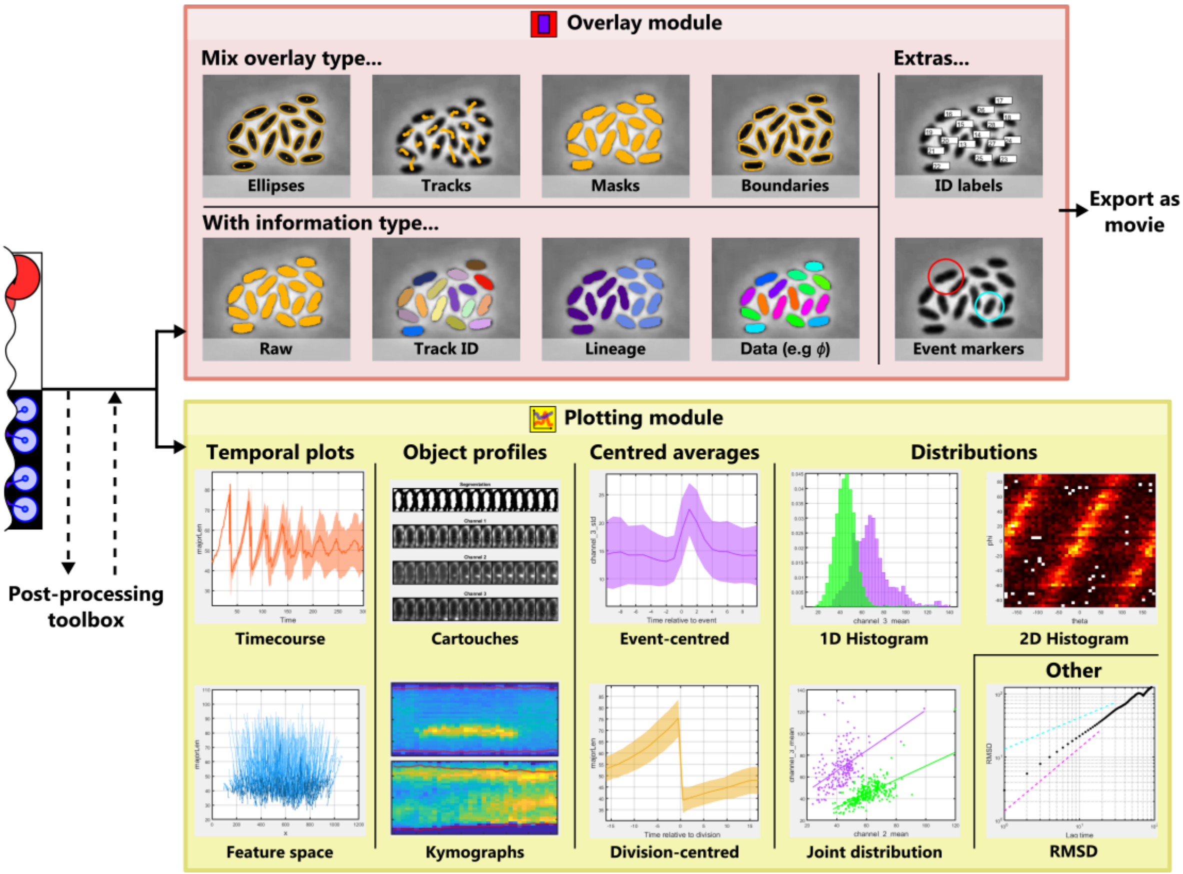

**Figure S5:** The visualisation modules. In both the case of the Overlays module (upper) and the Plotting module (lower), examples of each type of visualisation that can be achieved with the GUI are shown.

##### 3.6 Data visualisation modules

The visualisation modules allow rapid investigation of hypotheses and validation of analyses (figure S5). The Overlays module allows feature and track information to be presented directly on top of raw microscopy data, while the Plotting module provides GUI-based tools for generating a wide variety of different data visualisations from track data.

To use the Overlays module, users select:

1. The underlay: one of the available imaging channels or the derived segmentation.
2. The overlay type: **ellipses** - for elliptical reconstructions of objects, **tracks** - for plotting of trajectories, **masks** - for the segmentation, or **boundaries** - for the exterior of the segmentation.
3. The information type: **raw** - to colour all objects uniformly, **track ID** - to colour objects according to their corresponding track index, **lineage** - to colour objects according to their original mother cell, or **data** - to colour objects according to the value of a specified feature or associated quantity.

In addition to these options, users can apply track ID labels to the image, allowing the track data corresponding to visually identified objects of interest to be easily located. Once the overlay has been set up, users can apply the chosen settings to every frame of the dataset and automatically generate .tif images of the resulting visualisations. These can then be converted into timelapse videos through a variety of tools, such as FIJI [22].

The Plotting module likewise provides numerous visualisation options:

- **Timecourse** - Indicates how a specified feature varies over time.
- **Feature space** - Plots tracks on a user-specified pair of feature axes. Time is indicated by a gradient of shading along the track length, progressing from dark to light.
- **Cartouches** - Creates several timeseries of images for the specified track, with images cropped from the raw input from each channel.
- **Kymographs** - Generates space-time plots of a specified track from all channels. Similar to the 'Cartouches' option, but with each timepoint represented by a single 1D list of values.
- **Division-centred average** - feature timecourses aligned at the time of cell division and averaged across all divisions.
- **1D histogram** - A histogram of a specified feature. Users can choose whether to treat all feature values within a track as independent, or average them across the track.
- **Joint distribution** - A scatter plot comparing a specified pair of features.
- **2D histogram** - A heatmap indicating the joint distribution between a specified pair of features. Similar to the 'Joint distribution' option, but useful for large datasets where scatterplots are too crowded.
- **RMSD** - Root mean squared displacement. Displays the average distance an object moves from its starting position over a given lag time.

Once a plot has been generated, it can be exported as either a .tif bitmap or a .fig Matlab file. FAST also allows export of the raw data used to generate the figure, allowing users to easily access processed data for further analysis or to use in bespoke plots.

##### 3.6.1 Population and event labelling

Following processing by the Tracking and (optionally) the Division Detection modules, data can be annotated with population and event labels using the post-processing toolbox (`https://bit.ly/3Dbwst7`). These annotations are necessary for certain tools within the visualisation modules to be unlocked.

In the overlays module, events can be indicated as coloured circles around corresponding objects at the specified timepoint. Different classes of event can be overlaid using different colours.

Additionally, the following features become available within the plotting module:

- Addition of event labels allows plotting of event-centred averages, which are averaged timecourses aligned by a specific class of event.
- In the timecourse, phase space, event-centred average, division-centred average, 1D histogram, joint distribution and RMSD plotting options, track data can be split and coloured according to the population label assigned to each track. Some plotting options also support the application of statistical comparisons between different populations.

#### 3.7 Batch processing

FAST also includes a batch processing system, doubleFAST. To use this system, users simply need to analyse a single dataset using the standard FAST pipeline. doubleFAST will then read the settings files generated by this initial analysis, and automatically apply the same settings to all datasets provided.

### 4 Appendix A: Entropy of a uniformly sampled random vector

For a random vector $\boldsymbol{X}$ uniformly sampled from its support $\Omega$, the corresponding PDF $f(\boldsymbol{x})$ can be written as:

$$f(\boldsymbol{x}) = \begin{cases} (\int_\Omega 1 \cdot d\boldsymbol{x})^{-1} & \text{for} \quad \boldsymbol{x} \in \Omega \\ 0 & \text{for} \quad \boldsymbol{x} \notin \Omega. \end{cases} \tag{20}$$

From equation 10, we have:

$$\begin{aligned} H(\boldsymbol{X}) &= -\int_\Omega f(\boldsymbol{x}) \log_2(f(\boldsymbol{x})) \cdot d\boldsymbol{x}, \\ &= -\int_\Omega \left(\int_\Omega 1 \cdot d\boldsymbol{x}\right)^{-1} \log_2\left[\left(\int_\Omega 1 \cdot d\boldsymbol{x}\right)^{-1}\right] \cdot d\boldsymbol{x}, \\ &= -\left(\int_\Omega 1 \cdot d\boldsymbol{x}\right)^{-1} \log_2\left[\left(\int_\Omega 1 \cdot d\boldsymbol{x}\right)^{-1}\right] \int_\Omega 1 \cdot d\boldsymbol{x}, \\ &= \log_2\left(\int_\Omega 1 \cdot d\boldsymbol{x}\right). \end{aligned} \tag{21}$$

### References

[1] A. Ducret, E. M. Quardokus, and Y. V. Brun, "MicrobeJ, a tool for high throughput bacterial cell detection and quantitative analysis," *Nature Microbiology*, vol. 1, pp. 1–7, jun 2016.

[2] A. Paintdakhi, B. Parry, M. Campos, I. Irnov, J. Elf, I. Surovtsev, and C. Jacobs-Wagner, "Oufti: an integrated software package for high-accuracy, high-throughput quantitative microscopy analysis," *Molecular Microbiology*, vol. 99, pp. 767–777, feb 2016.

[3] S. Stylianidou, C. Brennan, S. B. Nissen, N. J. Kuwada, and P. A. Wiggins, "SuperSegger: robust image segmentation, analysis and lineage tracking of bacterial cells," *Molecular Microbiology*, vol. 102, pp. 690–700, nov 2016.

[4] J.-Y. Tinevez, N. Perry, J. Schindelin, G. M. Hoopes, G. D. Reynolds, E. Laplantine, S. Y. Bednarek, S. L. Shorte, and K. W. Eliceiri, "TrackMate: An open and extensible platform for single-particle tracking," *Methods*, vol. 115, pp. 80–90, feb 2017.

[5] Q. Wang, J. Niemi, C.-M. Tan, L. You, and M. West, "Image segmentation and dynamic lineage analysis in single-cell fluorescence microscopy," *Cytometry Part A*, vol. 77A, pp. 101–110, jan 2009.

[6] J. Klein, S. Leupold, I. Biegler, R. Biedendieck, R. Münch, and D. Jahn, "TLM-tracker: Software for cell segmentation, tracking and lineage analysis in time-lapse microscopy movies," *Bioinformatics*, vol. 28, pp. 2276–2277, sep 2012.

[7] R. Hartmann, M. C. F. van Teeseling, M. Thanbichler, and K. Drescher, "BacStalk: A comprehensive and interactive image analysis software tool for bacterial cell biology," *Molecular Microbiology*, vol. 114, pp. 140–150, apr 2020.

[8] D. J. Barry, C. H. Durkin, J. V. Abella, and M. Way, "Open source software for quantification of cell migration, protrusions, and fluorescence intensities," *The Journal of Cell Biology*, vol. 209, pp. 163–80, apr 2015.

[9] M. Linkert, C. T. Rueden, C. Allan, J. M. Burel, W. Moore, A. Patterson, B. Loranger, J. Moore, C. Neves, D. MacDonald, A. Tarkowska, C. Sticco, E. Hill, M. Rossner, K. W. Eliceiri, and J. R. Swedlow, "Metadata matters: Access to image data in the real world," *Journal of Cell Biology*, vol. 189, pp. 777–782, may 2010.

[10] T. Lindeberg, "Edge Detection and Ridge Detection with Automatic Scale Selection," *International Journal of Computer Vision*, vol. 30, no. 2, pp. 117–156, 1998.

[11] Y. Nishigami, M. Ichikawa, T. Kazama, R. Kobayashi, T. Shimmen, K. Yoshikawa, and S. Sonobe, "Reconstruction of Active Regular Motion in Amoeba Extract: Dynamic Cooperation between Sol and Gel States," *PLoS ONE*, vol. 8, aug 2013.

[12] F. Federici, "Arabidopsis thaliana plant cells containing chloroplasts, LM - CC BY 4.0," 2019. Available at: https://bit.ly/3gilcSH.

[13] R. Wheeler, "Human Erythrocytes Osmotic Pressure Phase Contrast Plain - CC BY-SA 3.0," 2012. Available at: https://bit.ly/3D5diUH.

[14] F. Meyer, "Topographic distance and watershed lines," *Signal Processing*, vol. 38, pp. 113–125, jul 1994.

[15] D. Geiger, A. Gupta, L. Costa, and J. Vlontzos, "Dynamic programming for detecting, tracking, and matching deformable contours," *IEEE Transactions on Pattern Analysis and Machine Intelligence*, vol. 17, pp. 294–302, mar 1995.

[16] A. Tsai, A. Yezzi, W. Wells, C. Tempany, D. Tucker, A. Fan, W. Grimson, and A. Willsky, "A shape-based approach to the segmentation of medical imagery using level sets," *IEEE Transactions on Medical Imaging*, vol. 22, pp. 137–154, feb 2003.

[17] S. Aksoy and R. M. Haralick, "Feature normalization and likelihood-based similarity measures for image retrieval," *Pattern Recognition Letters*, vol. 22, pp. 563–582, apr 2001.

[18] T. M. Cover and J. A. Thomas, *Elements of Information Theory*. Wiley Series in Telecommunications, New York, USA: John Wiley & Sons, Inc., 1st ed., 1991.

[19] N. A. Ahmed and D. V. Gokhale, "Entropy Expressions and Their Estimators for Multivariate Distributions," *IEEE Transactions on Information Theory*, vol. 35, no. 3, pp. 688–692, 1989.

[20] O. Al-Kofahi, R. J. Radke, S. K. Goderie, Q. Shen, S. Temple, and B. Roysam, "Automated Cell Lineage Construction: A Rapid Method to Analyze Clonal Development Established with Murine Neural Progenitor Cells," *Cell Cycle*, vol. 5, pp. 327–335, feb 2006.

[21] E. Meijering, O. Dzyubachyk, and I. Smal, "Methods for Cell and Particle Tracking," *Methods in Enzymology*, vol. 504, pp. 183–200, jan 2012.

[22] J. Schindelin, I. Arganda-Carreras, E. Frise, V. Kaynig, M. Longair, T. Pietzsch, S. Preibisch, C. Rueden, S. Saalfeld, B. Schmid, J.-Y. Tinevez, D. J. White, V. Hartenstein, K. Eliceiri, P. Tomancak, and A. Cardona, "Fiji: an open-source platform for biological-image analysis," *Nature Methods*, vol. 9, pp. 676–682, jul 2012.
